# Disruption of nucleolar integrity triggers cellular quiescence through organelle rewiring and secretion

**DOI:** 10.1101/2025.08.28.672879

**Authors:** Prashant Rawat, Tabea Quaderer, Ino Karemaker, Sung Sik Lee, Federico Ulliana, Zacharias Kontarakis, Jacob E. Corn, Matthias Peter

## Abstract

The nucleolus is the largest membraneless nuclear organelle and a critical regulator of growth and stress responses, comprised of over 600 proteins involved in ribosome biogenesis. However, how the nucleolar function and architecture coordinate cell-wide adaptive programs remains unclear. Here, we combined multi-omics profiling with functional genomics to define the cellular consequences of nucleolar disruption. We identify a stress response pathway connecting the **<u>nu</u>**cleolus to the **<u>G</u>**olgi apparatus, **<u>e</u>**ndo-lysosomal **t**rafficking, and cellular **<u>s</u>**ecretion (NuGETS). Chronic nucleolar defects activate a TP53-dependent transcriptional program that promotes organelle expansion, enhances secretory activity, and induces cell cycle exit with prolonged quiescence and partial epithelial-to-mesenchymal transition (pEMT) phenotypes. A genome-wide CRISPRi screen uncovered 400 regulators of secretory and quiescent states upon nucleolar stress. These pathways are enriched in cancer progression associated with ribosomopathies. Our findings therefore link nucleolar stress responses and cancer development, revealing cell-wide regulatory mechanisms that safeguard survival when nucleolar function is compromised.

## Introduction

Cellular stress responses to various intra- and extracellular conditions are vital for adaptation and organismal survival^1^. While extrinsic environmental challenges trigger a coordinated stress response across different cellular compartments^1,2^, intrinsic-derived stress such as misregulation of one organelle often results in a general loss of cellular homeostasis^3,4^. Cell-wide mapping of intrinsic stress responses revealed important insight into the function of membranous organelles^5^. In contrast, the stress regulation of membraneless organelles or biomolecular condensates remains largely unknown, in part due to the sub-microscopic dimensions and transient interactomes.

The largest membraneless structure inside the nucleus is the nucleolus, which is essential for cell growth, coordinating stress responses and cell survival^6–8^. The nucleolus orchestrates the first steps of ribosome biogenesis, and in metazoans, the distinct phases are spatially organized within nucleolar sub-compartments^9,10^. The rDNA transcription by RNA Polymerase I (Pol I) occurs in the innermost fibrillar centre (FC) followed by rRNA processing in the middle dense fibrillar component (DFC), and finally assembly of pre-ribosomal particles in the outermost granular component (GC)^9,11^. Recent work highlighted biochemical and cellular principles of how nucleolar sub-compartments are formed and maintained^12–15^. Moreover, novel mapping tools revealed how nucleolar architecture remodels in response to defects in distinct steps of ribosome biogenesis^16^.

Given the link between architecture, physical properties and its function, the nucleolus provides an attractive model to study structure-function relationships within a biomolecular condensate. Indeed, previous studies have correlated nucleolar morphology and associated changes with the loss of nucleolar components^17,18^, loss of heterochromatin anchoring^19,20^, loss of nucleolar associated domains (NADs)^21,22^, premature ageing^23^, and reported hallmarks of neurodegeneration^24–26^ and cancer^27^. In fact, changes in nucleolar number and shape are often used as clinical diagnostic markers for many cancer types^28^. Moreover, hypo-sufficiency of nucleolar and ribosomal proteins leads to developmental abnormalities and promotes cancer progression^29,30^. Thus, defects in nucleolar architecture and function are associated with many human pathologies, underscoring the importance of dissecting the effects of chronic nucleolar misregulation on cellular homeostasis and disease progression^16,31^.

Although the long-term consequences of nucleolar alterations are critically important, studies to-date typically focused on nucleolar changes triggered by acute stress conditions such as heat shock or treatment with pharmacological agents^32–34^. Different nucleolar morphologies are reported to be associated with acute stresses, including formation of nucleolar caps upon RNA Pol I inhibition^35^ or of a “bare” rDNA scaffold upon transcriptional CDK inhibition^33^. Upon acute stress, ribosome biogenesis and rRNA processing is rapidly halted and the nucleolus serves as a buffering compartment for misfolded proteins^34,36,37^, facilitating protein refolding during recovery^38,39^. Moreover, ribosome biogenesis defects are thought to imbalance protein translation and increase mistranslated proteins, thereby activating cytoplasmic quality control pathways^38^.

Inhibition of different steps in ribosome biogenesis prevents integration of ribosomal protein (RP) subunits into the pre-ribosomal particles (SSU/LSU)^40^, triggering nucleolar stress. These free RP subunits, and in particular RPL5 and RPL11, bind MDM2/MDM4, the E3 ubiquitin ligases that target TP53 for proteasomal degradation. RP binding antagonizes MDM2/MDM4 activity, hence stabilizing TP53 levels^41–43^. TP53 is a potent transcription factor, which initiates a transcriptional program orchestrating cell proliferation and apoptosis by upregulating targets such as CDKN1A^44^ or the BH3 proteins PUMA and NOXA^45^. Selective depletion of RP subunits was also shown to induce a nucleolar phenotype. Of note, RPL5 or RPL11 depletion leads to distinct nucleolar morphology, and this defect is associated with increased DNA damage and cellular senescence^46^. Moreover, mutations in RPL5 are found in 28% of melanoma and 34% of breast cancer samples^47^.

Although previous studies investigated how loss of nucleolar proteins affect nucleolar morphology and translational phenotypes^48,49^, the cellular stress responses triggered by prolonged defects in nucleolar architecture and ribosome biogenesis remained unclear. To close this gap, we depleted structural proteins from different nucleolar compartments and examined their impact on cell physiology using an unbiased systems approach. Specifically, we RNAi-depleted the upstream binding transcription factor (UBF) from the FC, the U3 small nucleolar RNA-associated protein 18 homolog (UTP18) and Fibrillarin (FBL) from the DFC, and Nucleophosmin 1 (NPM1) from the GC. Of note, many nucleolar proteins such as FBL are essential and embryonically lethal, whereas their depletion in differentiated cells show minimal defects^50–52^. FBL is a conserved rRNA methyltransferase that associates with small nuclear RNAs, and defects in rRNA methylation alters rRNA processing with implications for DFC morphology^52,53^. UTP18 being part of the UTP-B complex also resides in the DFC, binds 5’ETS and supports rRNA processing^8^, while NPM is structurally essential for maintaining the GC and chromatin tethering^14,21^.

We discovered a distinct stress response pathway allowing cells to adapt to the loss of nucleolar architecture and ribosome biogenesis defects. These adaptation mechanisms require early transcriptional reprogramming and cytoplasmic organellar reorganization leading to partial EMT phenotypes and thus contribute to progression of different cancer types characteristic for late stages of ribosomopathies.

## Results

### Loss of nucleolar proteins leads to cell proliferation defects, nucleolar reorganization and ribosome biogenesis defects

To examine the role of nucleolar proteins in cell growth and proliferation, we siRNA-treated RPE1 cells to individually deplete NPM, UBF, and UTP18 and FBL proteins from the nucleolar GC, FC and DFC regions, respectively. Cell growth was monitored with MTT assays over a period of days. While we observed no proliferation defect after NPM depletion, proliferation significantly decreased after siRNA-targeting UBF and UTP18 and almost stopped in cells lacking FBL **(Fig 1A and S1A)**. FBL-depleted cells exhibited prolonged proliferation defects showing minimal cell growth in MTT assays up to 192 to 240 hours post siRNA transfections with effects peaking already at 96h **(Fig S1B)**. Live-cell imaging of FUCCI-RPE1 cells for 72h revealed a G1 cell cycle arrest post 30h siRNA treatment **(Fig 1B and Fig S1C-D)**. Specific depletion of the targeted nucleolar proteins was confirmed by immunofluorescence microscopy (**Fig 1C and Fig S1E)**. Interestingly, confocal microscopy at 96h post siRNA transfection revealed that FBL depletion led to dramatic nucleolar reorganization, resulting in a single large spherical nucleolus as compared to three to five irregularly shaped nucleoli in control cells. Similarly, depletion of UBF or UTP18 generally reduced the number of nucleoli and caused cells to establish one nucleolus. The nucleoli in cells lacking UBF were smaller, resembling the phenotype associated with decreased RNA Pol I activity^34^, whereas UTP18 depleted cells showed more irregularly shaped nucleoli **(Fig 1C-D and S1F)**. The single nucleolus formed upon FBL depletion still contained nucleolar proteins such as UBF, UTP18 and NNP1 **(Fig S1E and G)**. At longer time points, proliferation was restored due to FBL re-expression, and cells could reestablish the distinct nucleolar compartments, indicating that cells remain viable without FBL and with transient loss of nucleolar integrity **(Fig S1H)**.

**Figure 1:**
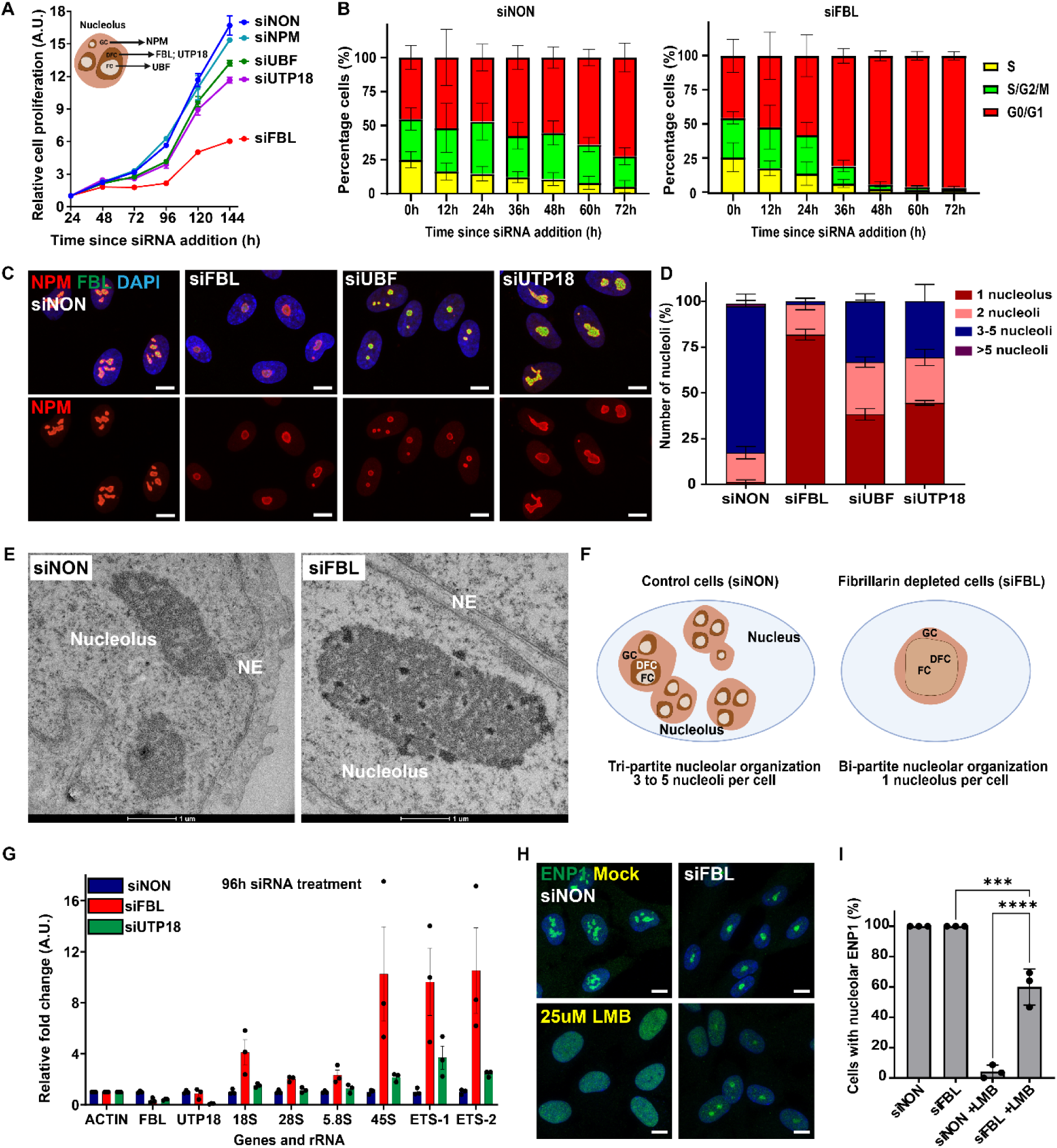
Depletion of nucleolar proteins leads to prolonged cell cycle arrest, loss of nucleolar architecture and ribosome biogenesis defects. **A.** Line graph shows cell proliferation determined by MTT assay of RPE1 cells over time in hours (h) relative to addition of siRNA against non-targeting control (siNON) or upon siRNA depletion of the indicated proteins localizing to different sub-nucleolar compartments (inset schematic), including Nucleophosmin 1 (siNPM), Upstream binding factor 1 (siUBF), U3 small nucleolar RNA-associated protein 18 homolog (siUTP18) and Fibrillarin (siFBL). Measurements are from N=4 samples. The 24-hour siRNA value was set to 1, and all other values were normalized to the 24-hour time point. **B.** Bar graph plotting the percentage (%) of cells in different cell cycle phases (S = yellow; S/G2/M = green; G0/G1=red) during live-cell imaging using FUCCI-RPE1 cells treated with siNON (left panel) or siFBL (right panel) at the indicated time in hours (h) relative to siRNA addition. Mean with SD is plotted from N=4 biological replicates. **C.** Representative confocal microscopic images showing RPE1 cells following treatment with the indicated siRNA for 96 hours (siNON, siFBL, siUBF, siUTP18), with the nucleolus/nucleus visualized by co-staining of NPM (red), FBL (green) and DAPI (blue). The top row displays the merged co-localization, and the bottom row NPM localization. Images are representative of N=4 biological replicates. Scale bar: 10 µm **D.** Bar graph showing the percentage (%) of RPE1 cells classified as having 1 (maroon), 2 (pink), 3-5 (navy blue) or >5 (purple) nucleoli, as observed with NPM staining. Mean with SEM is plotted from N=4 biological replicates. **E.** Transmission electron micrographs show nucleolar architecture of RPE1 cells upon siNON and siFBL at 96 hours siRNA treatment. The nucleolus and nuclear envelope (NE) are indicated. Scale bar: 1 µm. **F.** Schematic diagrams depicting nucleolar architecture observed in control (siNON; left panel) and FBL-depleted RPE1 cells (siFBL; right panel). **G.** Bar graph showing the fold change of transcripts corresponding to the indicated genes and ribosomal RNA species detected by RT-qPCR performed on cDNA derived from siNON-, siFBL- and siUTP18-depleted RPE1 cells at 96 hours post siRNA treatment. Actin RNA levels in siNON treated cells were set to 1, and all other values were normalized. Mean with SD is plotted from N=3 biological replicates. **H.** Representative confocal microscopic images of RPE1 cells treated with siNON and siFBL for 96 hours, at which point 25nM Leptomycin (LMB) or solvent control (mock) was added for 2 hours. Samples were co-stained for ENP1 (green) and DAPI (blue). Images are representative of N=3 biological replicates. Scale bar: 10 µm. **I.** Bar graph of data from (H) with RPE1 cells scoring cells with nucleolar ENP1 signal. Mean with SD is plotted from N=3 biological replicates. Statistical analysis was performed with an ordinary one-way Anova test with Tukey’s multiple comparison.

To better visualize the nucleolar morphology, we compared transmission electron microscopy (TEM) images of RPE1 cells treated with siFBL or non-targeting control (siNON) oligos. While nucleoli in control cells appeared as dense nuclear structures, with less dense FC surrounded by denser DFC/GC regions, the rearranged nucleolus in FBL-depleted cells accumulated very dense patches surrounding lighter FC regions **(Fig 1E)**, indicating rRNA processing defects. Increased nucleolar density in FBL-depleted cells was also visible by label-free quantitative phase imaging (QPI) microscopy (**Fig S1I-J)**, excluding fixation artifacts. Surprisingly, we observed a significant loss of nucleoplasm density in FBL-depleted cells, indicating a loss of heterochromatin upon nucleolar reorganization **(Fig S1K)**. Altogether, we conclude that depletion of different nucleolar components in RPE1 cells results in distinct nucleolar morphologies **(Fig 1F)**, including a predominant one-nucleolar phenotype associated with a cell cycle delay in G1.

To test whether these nucleolar morphology changes cause rRNA processing defects, we quantified the abundance of various rRNA species by RT-qPCR in RPE1 cells that were siRNA-depleted of FBL or UTP18. While mRNA levels of the targeted components were reduced as expected, FBL transcripts were also lost in UTP18-depleted cells at late time points **(Fig 1G)**. Conversely, UTP18 mRNA levels were unaffected in FBL-depleted cells. Importantly, 45S, ETS1 and ETS2 species accumulated after siRNA depletion of either FBL or UTP18 **(Fig 1G and Fig S1L)**. These early rRNA processing defects appeared constant in UTP18-depleted cells during the entire time course, while the defects became more pronounced in FBL-depleted cells over time. Additionally, FBL-depleted cells showed significantly higher accumulation of 18S and 28S rRNA species **(Fig 1G and Fig S1L)**, confirming defects in early rRNA processing. To corroborate these data, we followed the localization of the rRNA processing factor ENP1 (BYSL) as a marker for ribosome biogenesis^54^. As expected, ENP1 was localized in the nucleolus and nucleoplasm in both control- and fibrillarin-depleted cells. However, after inhibiting nuclear export of pre-ribosomal particles by Leptomycin B (LMB), ENP1 was retained in the nucleoplasm in the control (siNON) but not in FBL-depleted cells, which showed nucleolar ENP1 **(Fig 1H-I)**. Together, these data suggest that cells lacking FBL exhibit a ribosome biogenesis defect, most likely because of aberrant rRNA processing in the nucleolus and subsequent nucleolar reorganization.

### Persistent nucleolar defects only mildly perturb Pol I and Pol2 transcription, without decreasing the levels of cytoplasmic ribosomes and protein translation

To measure changes in total RNA Pol II activity, we probed for nascent transcription using the uridine analogue 5-EU with short and long incubations and observed sustained transcription despite severe nucleolar and cell cycle defects. In addition, the nascent RNA signal accumulated in the nucleolus with longer incubations, confirming active transcription of both RNA Pol I and RNA Pol II **(Fig S2A-D)**. To further corroborate these data, we probed for Ser2 and Ser5 phosphorylation marks indicative of active transcription. While total RNA Pol II levels in FBL-depleted cells slightly decreased, Ser2P and Ser5P showed significantly higher levels **(Fig 2A-D)**, indicating ongoing and active RNA Pol II transcription. Similarly, total RNA Pol II levels mildly decreased in cells lacking UBF, while Ser5P was upregulated with no changes in Ser2P levels. In contrast, UTP18-depleted cells showed no significant changes in total RNA Pol II and Ser2P but significantly reduced Ser5P levels **(Fig 2B-D and Fig S2E)**. These patterns suggest active RNA Pol II transcription with widespread pausing in the absence of FBL and UBF, and defects in transcription initiation and promoter proximal pausing in the absence of UTP18. We conclude that RNA Pol II remains active in cells exposed to persistent nucleolar defects induced by depleting various nucleolar proteins, with specific transcription pausing and initiation defects.

**Figure 2:**
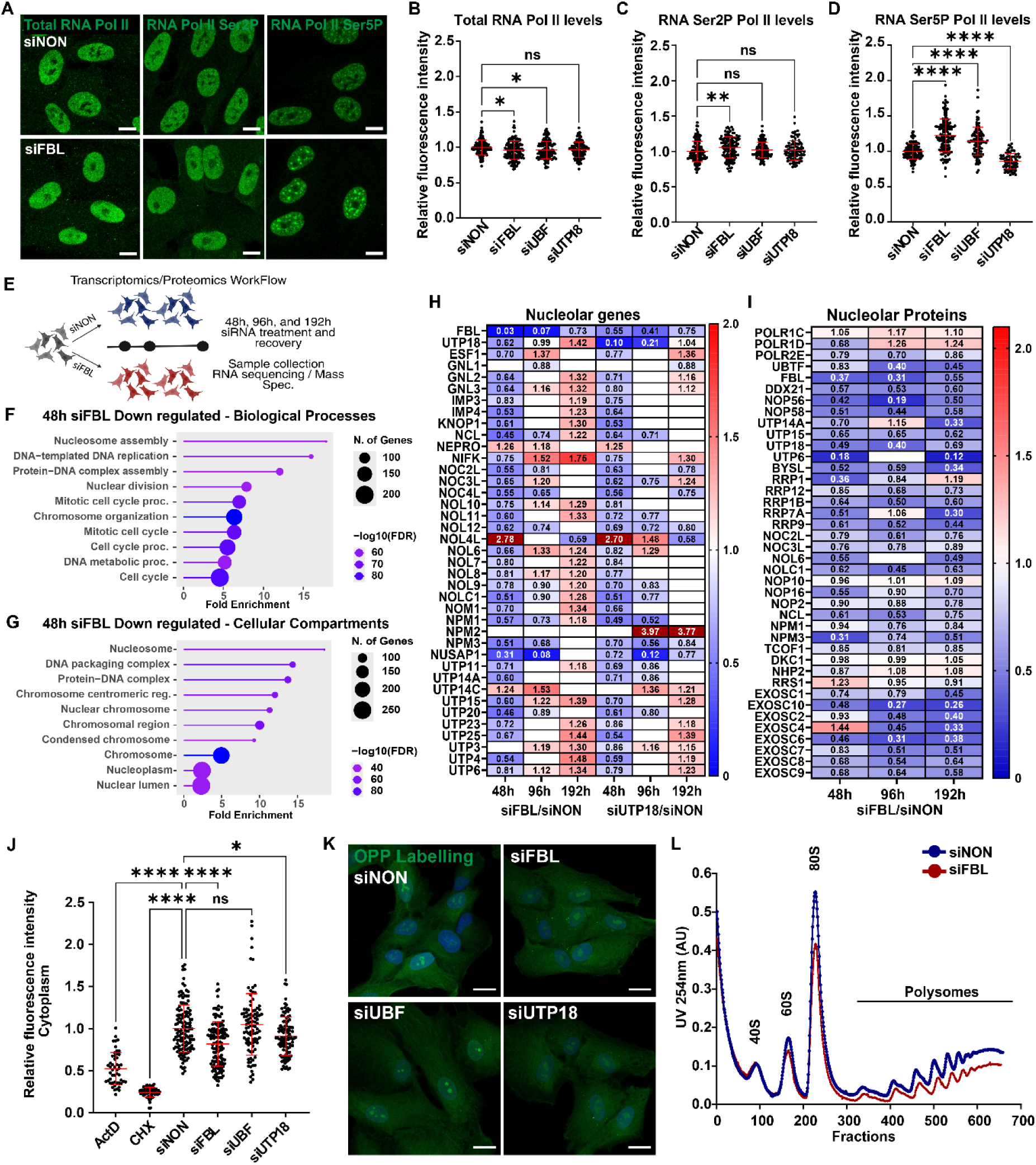
Chronic nucleolar defects lead to selective downregulation of nucleolar genes without affecting total gene-expression. **A.** Representative confocal microscopic images showing nuclei of RPE1 cells treated with siNON (top row) or siFBL (bottom row) for 96 hours, and stained for total RNA Polymerase II (green, left), serine-2-phosphorylation of the C-terminal tail of RNA Polymerase II (green, middle) and serine-5-phosphorylation of the C-terminal tail of RNA polymerase II (green, right). Images are representative of N=3 biological replicates. Scale bar: 10 µm. **B.** Dot plot showing data from Fig 2A and S2E, measuring the fluorescence intensity of total RNA Polymerase II in the nucleus of RPE1 cells treated as indicated with siRNA for 96 hours (siNON, siFBL, siUBF, siUTP18). All values were normalized to the siNON average of the corresponding experiment. Mean with SD is plotted from N=3 biological replicates. Statistical analysis was performed by an ordinary one-way Anova test with Dunnett’s multiple comparison. Asterisks denote p value (*, <0.05); ns, non-significant. **C.** Dot plot showing data from Fig 2A and S2E, measuring the nuclear fluorescence intensity of serine-2-phosphorylation of the C-terminal tail of RNA Polymerase II following treatment with the indicated siRNA for 96 hours (siNON, siFBL, siUBF, siUTP18). All values were normalized to siNON average of the corresponding experiment. The mean with SD is plotted from N=3 biological replicates. Statistical analysis was performed by an ordinary one-way Anova test with Dunnett’s multiple comparison. Asterisks denote p value (**, <0.01). ns, non-significant. **D.** Dot plot showing data from Fig 2A and S2E, measuring the nuclear fluorescence intensity of serine-5-phosphorylation of the C-terminal tail of RNA Polymerase II following treatment with the indicated siRNA for 96 hours (siNON, siFBL, siUBF, siUTP18). All values were normalized to siNON average of the corresponding experiment. The mean with SD is plotted from N=3 biological replicates. Statistical analysis was performed by an ordinary one-way Anova test with Dunnett’s multiple comparison. Asterisks denote p value of (****, <0.0001). ns, non-significant. **E.** Schematic diagram depicting the experimental workflow and time-points for the transcriptomics (RNA-seq) and proteomic (mass-spectrometry) analysis. **F.** Lollipop chart of biological processes enrichment analysis using ShinyGO from the RNA-seq dataset for genes with more than 1.5-fold downregulation and a p-value cutoff of 0.05 of RPE1 cells treated with siFBL or siNON siRNA oligos for 48 hours. Pathways over fold enrichment were plotted with -log10FDR and represented using the indicated color scheme. Number of genes: 528 and FDR cutoff: 0.01. **G.** Lollipop chart of cellular component enrichment analysis using ShinyGO from RNA-seq dataset for genes with more than 1.5-fold downregulation and a p-value cutoff of 0.05 of RPE1 cells treated with siFBL or siNON siRNA oligos for 48 hours. Pathways over fold enrichment were plotted with -log10FDR and represented using the indicated color scheme. Number of genes: 528 and FDR cutoff: 0.01. **H.** Heat map representing gene fold change of nucleolar proteins from RNA-seq data of RPE1 cells treated as indicated with siFBL, siUTP18 or siNON controls for 48-, 96- and 192 hours. The scale bar ranges from 0 to 2-fold change and genes follow a p-value cutoff of 0.05. Out of scale values are represented in maroon. Non-significant values are left blank and represented in white. The gene list was assembled based on the available literature. **I.** Heat map representing protein level fold change of nucleolar proteins from the total cellular proteome measured by mass spectrometry of RPE1 cells treated with siFBL or siNON oligos for 48-, 96- and 192 hours. The scale bar ranges from 0 to 2-fold change and proteins follow a p-value cutoff of 0.05. Out of scale values are represented in maroon. Non-significant values are left blank and represented in white. The protein list was assembled based on the available literature. **J.** Dot plot data from Fig 2K and S2N, measuring the fluorescence intensity for O-Propargyl-puromycin with AF488-Azide signal in the cytoplasm of RPE1 cells treated with the indicated siRNA for 96 hours (siNON, siFBL, siUBF, siUTP18). For control, cells were exposed to 625nM Actinomycin D (ActD) or 30µg/ml Cycloheximide (CHX) for 2 hours. All values were normalized to the siNON average of the corresponding experiment. The mean with SD is plotted from N=3 (siRNA) and N=2 (inhibitors) biological replicates. Statistical analysis was performed by an ordinary one-way Anova test with Dunnett’s multiple comparison. Asterisks denote p value of (****, <0.0001; *, <0.05). ns, non-significant. **K.** Representative confocal microscopic images showing RPE1 cells treated with the indicated siRNA for 96 hours (siNON, siFBL, siUBF, siUTP18), and co-stained for O-Propargyl-puromycin (OPP) with AF488-Azide (green) and DAPI (blue). 10 µM OPP was added for 3 hours. Images are representative of N=3 biological replicates. Scale bar: 10 µm. **L.** Graph showing RNA absorbance at 260 nm measured along sucrose polysome gradients fractionating cell extracts prepared from RPE1 cells treated with siNON or siFBL for 96 hours. Graphs are representative of N=2 biological replicates.

To understand how cells rewire their gene expression upon nucleolar perturbation, we performed time-course omics experiments on FBL- and UTP18-depleted RPE1 cells. First, we compared steady state RNA and protein levels at three time points during siRNA treatment, based on cell proliferation data corresponding to an early (48h) or late stress response (96h), and at the onset of stress recovery (192h) **(Fig S1B, 2E and S2F)**. As expected, many downregulated genes and corresponding proteins are associated with cell cycle, mitosis, nucleosome assembly and DNA replication **(Fig 2F-G and S2G-I)**. Moreover, consistent with ribosome biogenesis defects, we measured a significant transcriptional downregulation of nucleolar components in both FBL- and UTP18-depleted cells **(Fig 2H-I)**, correlating with the strongest effects seen for rRNA processing like NOP and UTP proteins and structural components such as NCL and NPM. Interestingly, the RNA Pol I machinery remained unaltered at the protein level at early time points of nucleolar stress **(Fig 2I)**. Together, these results suggest that nucleolar defects trigger an early, coordinated transcriptional response that alters the nucleolar proteome and function.

Surprisingly, despite the ribosome biogenesis defects, ribosomal proteins showed minimal downregulation of both the transcriptome and proteome at 48h after FBL depletion, preserving more than 60% of the ribosomal proteins even at the 96h and 192h time points **(Fig S2J-K)**. Upon closer analysis, we noticed a discrepancy in expression of ribosomal biogenesis genes and ribosomal protein genes, both of which are transcribed by RNA Pol II. While ribosome biogenesis genes show a remarkable downregulation, genes encoding ribosomal proteins remain largely unaffected, indicating differential transcriptional regulation and feedback for these two distinct gene classes **(Fig 2H and S2J)**. Indeed, immunofluorescence analysis confirmed that levels of cytoplasmic ribosomal subunits remained constant **(Fig S2L-M)**, implying that nucleolar defects affect nucleolar proteins but do not greatly change the levels of cytoplasmic ribosomes. We conclude that the ribosomal pool has vast buffering capacity even in the absence of ribosome biogenesis, maintaining sufficient protein translation to preserve cell viability. To corroborate these findings, we monitored global protein translation using *O*-propargyl-puromycin (OPP), a cell permeable compound that through click chemistry can be covalently coupled to a fluorescent dye to visualize nascent proteins. As expected, almost no nascent protein labelling was detected when protein translation was blocked by cycloheximide (CHX) treatment, and protein translation was reduced significantly in cells treated with actinomycin (ActD) to block RNA polymerase activity **(Fig 2J and S2N-O)**. In UBF-depleted cells, the OPP staining showed no significant reduction in nuclear and cytoplasmic fractions but slightly increased levels in the nucleolus, possibly because of the reduced nucleolar size. Interestingly, FBL- and UTP18-depletion slightly reduced nascent protein levels in the nucleolar, nuclear and cytoplasmic compartments, **(Fig 2J-K and Fig S2O-P)**, but protein translation remained substantial despite the aberrant one-nucleolar morphology. Consistent with these results, polysome profiling of FBL-depleted cells confirmed reduced monosome (80S) and polysome fractions compared to controls, yet the cells preserve significant amounts of these species **(Fig 2L)**. We conclude that despite reduced Pol I and II transcription and ribosome biogenesis defects, cells with aberrant nucleolar morphology remain surprisingly robust and sustain sufficient protein translation to maintain essential cellular processes.

### Chronic nucleolar defects activate TP53 stress signaling

Interestingly, RNA-Seq analysis of FBL- and UTP18-depleted RPE1 cells revealed significant bidirectional changes in gene expression already at the earliest time point measured after siRNA treatment, indicating early transcriptional reprogramming **(Fig S3A-B)**. A similar bidirectional stress response signature was also apparent by proteomic analysis in FBL-depleted cells, albeit with delayed dynamics compared to transcriptomics changes and sustained proteome changes until the early recovery time point **(Fig S3C)**. TP53 is well-studied as an early sensor of nucleolar defects, genome instability and ribotoxic stress responses and its activation is linked to many downstream consequences such as DNA-damage repair, cell cycle arrest and apoptosis. Indeed, we observed a strong TP53 signature in the gene enrichment analysis of FBL and UTP18 knockdowns **(Fig 3A, Fig S3D-E)**, with robust upregulation of many TP53 targets including CDKN1A, TP53INP1 and TIGAR **(Fig 3B-C)**. As expected, immunofluorescence and western blotting confirmed TP53 stabilization and increased CDKN1A protein levels upon chronic nucleolar defects originating from depletion of nucleolar proteins **(Fig 3D-F, S3F-G)**. To determine whether TP53 activation is responsible for cell cycle arrest and nucleolar reorganization, we simultaneously siRNA-depleted FBL and TP53, and performed live cell FUCCI-RPE1 imaging and nucleolar staining. Indeed, co-depletion of TP53 or CDKN1A with FBL rescued cell cycle arrest without restoring FBL levels **(Fig 3F-H, S3H)**, and surprisingly also abrogated the one-nucleolar phenotype **(Fig 3I, S3I-K)**. Similarly, co-depletion of TP53 with UBF or UTP18 in RPE1 cells abrogated the one-nucleolus phenotype **(Fig S3 I-K)**. Overall, these results suggest that TP53 is activated by nucleolar defects and is in part responsible for cell cycle arrest and nucleolar reorganization.

**Figure 3:**
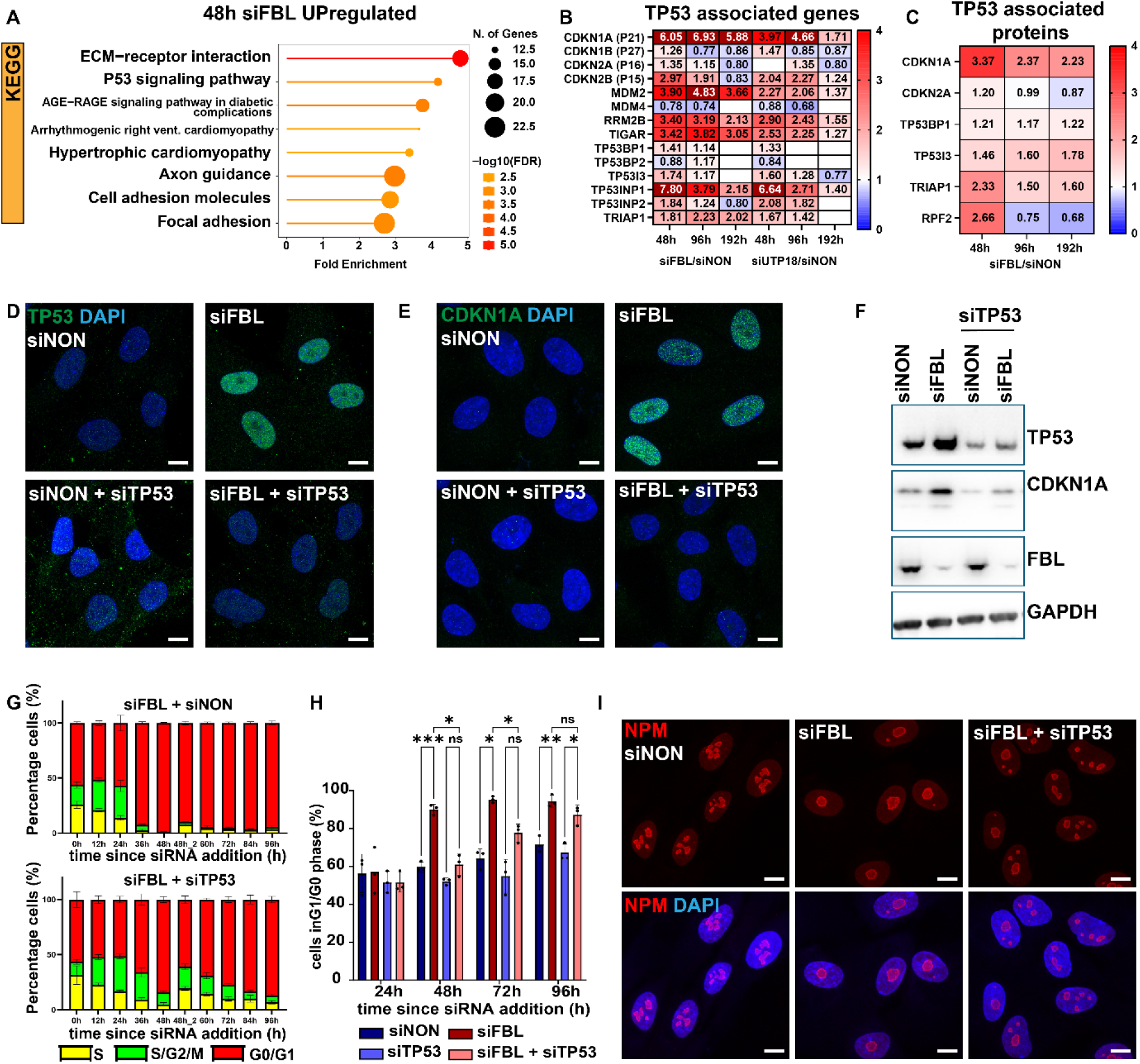
Prolonged nucleolar defects activate TP53 signaling. **A.** Lollipop chart of KEGG (Kyoto encyclopedia of genes and genomes) enrichment analysis from RNA-seq datasets prepared from RPE1 cells treated for 48 hours with siFBL or siNON controls using ShinyGO for genes with more than 2-fold upregulation and a p-value cutoff of 0.05. Pathways over fold enrichment were plotted with -log10FDR using the indicated color scheme. Number of genes: 978 and FDR cutoff: 0.01. **B.** Heat map representing genes fold change of TP53-associated proteins from RNA-seq datasets of RPE1 cells treated for 48-, 96- and 192 hours with siFBL, siUTP18 and siNON controls. The scale bar ranges from 0 to 4-fold change and genes follow a p-value cutoff of 0.05. Out of scale values are represented in maroon. Non-significant values are left blank and represented in white. The gene list is based on the available literature. **C.** Heat map representing protein level fold change of TP53 associated proteins from mass spectrometry measurements of the total cellular proteome, prepared from RPE1 cells treated for 48-, 96- and 192 hours with siFBL or siNON controls. The scale bar ranges from 0 to 4-fold change and proteins follow a p-value cutoff of 0.05. The protein list is based on available literature. **D.** Representative confocal microscopic images showing RPE1 cells treated with the indicated single or double siRNAs for 96 hours (siNON, siFBL, siNON+siTP53, siFBL+siTP53), and co-stained for TP53 (green) and DAPI (blue). Images are representative of N=4 biological replicates. Scale bar: 10 µm. **E.** Representative confocal microscopic images showing RPE1 cells treated with the indicated single or double siRNAs for 96 hours (siNON, siFBL, siNON+siTP53, siFBL+siTP53), and co-stained for CDKN1A (green) and DAPI (blue). Images are representative of N=4 biological replicates. Scale bar: 10 µm. **F.** Immunoblots of total cell lysates prepared from RPE1 cells treated with the indicated single or double siRNAs for 96 hours (siNON, siFBL, siNON+siTP53, siFBL+siTP53), and probed with antibodies against TP53, CDKN1A and FBL. GAPDH controls equal loading. **G.** Bar graph showing the percentage (%) of FUCCI-RPE1 cells in the different cell cycle phases (S = yellow; S/G2/M = green; G0/G1=red) using live-cell imaging of double knockdowns of siFBL + siNON (top panel) or siFBL + siTP53 (bottom panel) at the indicated time in hours (h) relative to siRNA addition. The cells were freshly re-plated to avoid confluency at 48h indicated as 48_2. The mean with SD is plotted from N=3 biological replicates. **H.** Bar graph showing data from Fig 3G and S3H, measuring the percentage (%) of FUCCI-RPE1 cells exposed to the indicated double knockdowns (siNON + siNON, siFBL + siNON, siNON + siTP53 or siFBL + siTP53) in the G0/G1 cell cycle phase at the indicated time points after siRNA addition. 48_2 was used for quantifications. The mean with SD is plotted from N=3 biological replicates. Statistical analysis was performed by a two-way Anova test with Tukey’s multiple comparison. Asterisks denote p value (***, <0.001; **, 0.01; *, 0.05). ns, non-significant. **I.** Representative confocal microscopic images showing RPE1 cells treated with the indicated single or double siRNAs for 96 hours (siNON, siFBL, siFBL+siTP53), and co-stained for NPM (red) and DAPI (blue). The top row displays NPM localization and the bottom row merged co-localization images. Images are representative of N=3 biological replicates (N=3). Scale bar: 10 µm.

### Persistent nucleolar defects trigger pronounced cytoplasmic rearrangements

Interestingly, Gene Ontology (GO) and KEGG pathway analysis from transcriptomics revealed that the depletion of nucleolar components strongly upregulates proteins associated with cytoskeletal organization, focal adhesion, the cell surface, and the extra cellular matrix (ECM) **(Fig 3A and S3D-E)**. Likewise, analysis of biological processes and cellular compartments confirmed striking enrichment of components of the ECM, the cell surface, the Golgi apparatus, the Golgi membrane, the lysosomal lumen and endosomal vesicles **(Fig 4A and S4A-D**), indicating that nucleolar defects increase expression of many cytoplasmic components. For example, the Golgi scaffolding and stacking regulators GOLGAs, COGs and GORASPs were upregulated in RNA-seq and proteomic experiments in both FBL- and UTP18-depleted cells **(Fig 4B and C)**. Increased expression of the Golgi marker GM130 (GOLGA2) upon depletion of all three nucleolar proteins was further validated by western blotting **(Fig 4D)** and confocal imaging **(Fig 4E-F)**. Importantly, upregulation of these components dramatically changed the morphology and size of the Golgi and other cytoplasmic membranous compartments. For example, immunofluorescence microscopy of GM130, GORASP1 and GORASP2 showed an enlarged Golgi apparatus with significant vasculo-tubular expansion in cells siRNA-depleted for either FBL, UBF or UTP18 **(Fig S4E-J)**. This Golgi expansion persisted in more than 30% of the cells even at 192h after siRNA addition **(Fig S4K)**. Similarly, nucleolar defects triggered upregulation of endosomal and lysosomal components **(Fig 4G-H)**, resulting in more and bigger EEA1 stained endosomes and TFEB- and LAMP1 marked endo-lysosomes **(Fig 4I-K)**. These experiments suggest that nucleolar defects activate TP53, which in turn not only alters nucleolar morphology, but also leads to a striking expansion of cytoplasmic organelles. Indeed, kinetic analysis indicates that nucleolar reorganization precedes cytoplasmic expansion of the Golgi apparatus and vesicular structures **(Fig 4L-M)**. Moreover, the enlargement of trafficking organelles was reversed when TP53 was co-depleted with FBL, UBF or UTP18 **(Fig S4 E-I and L-P)**, implying that the cytoplasmic reorganization is part of a TP53 response triggered by nucleolar defects.

**Figure 4:**
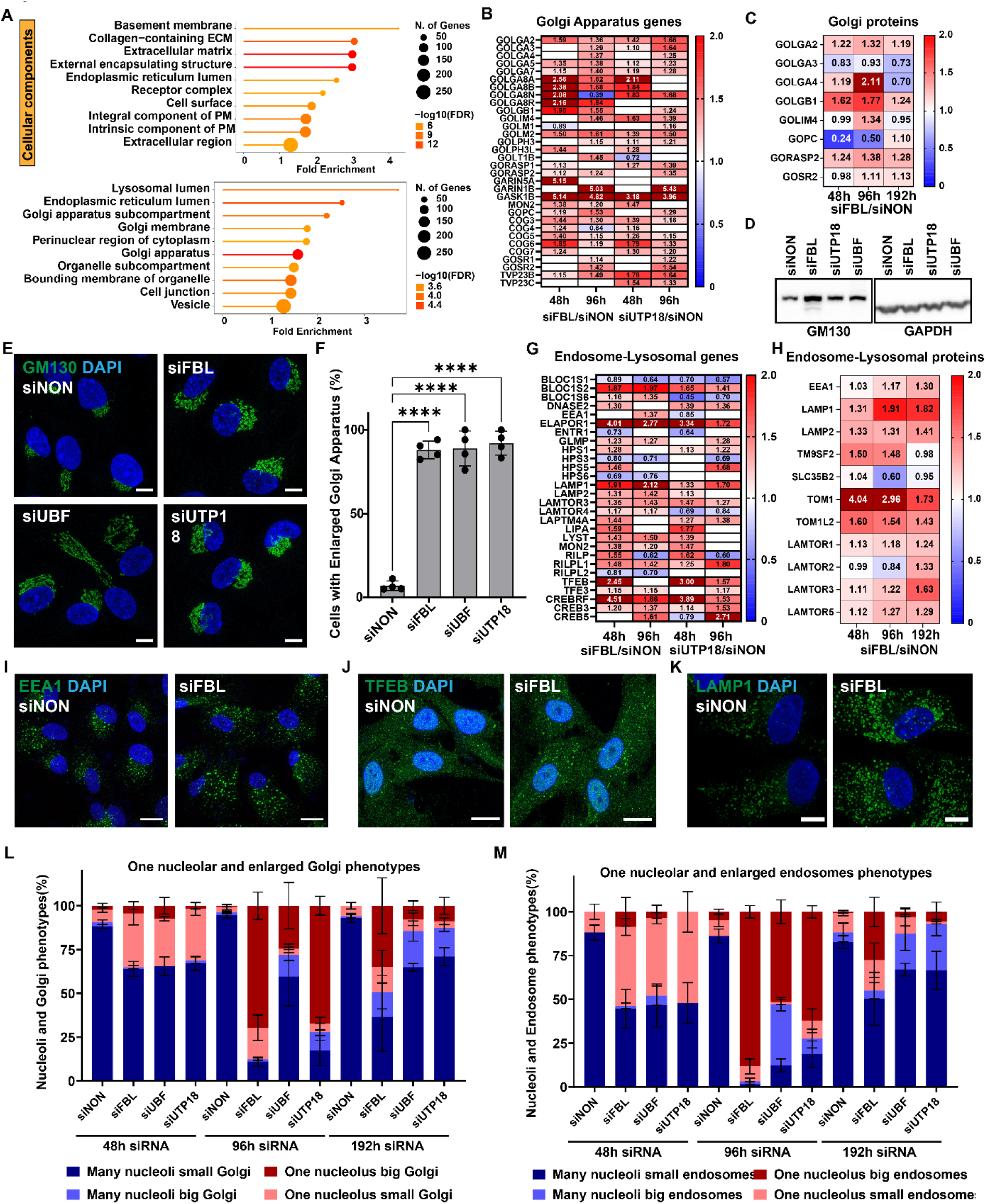
Chronic nucleolar defects induce prominent cytoplasmic rearrangements. **A.** Lollipop chart of cellular component enrichment analysis using RNA-seq datasets from Fig S3A prepared from RPE1 cells treated with siFBL or siNON controls for 48 hours using ShinyGO for genes with more than 2-fold (top panel) and 1.5-fold (bottom panel) upregulation and a p-value cutoff of 0.05. Pathways over fold enrichment were plotted with log10FDR using the indicated color scheme. Number of genes: 978 and 1077 respectively and FDR cutoff: 0.01. **B.** Heat map representing genes fold change of Golgi apparatus structural and functional components from RNA-seq dataset from Fig S3B generated from RPE1 cells treated with siFBL, siUTP18 or siNON controls for 48- and 96 hours. The scale bar ranges from 0 to 2-fold change and genes follow a p-value cutoff of 0.05. Out of scale values are represented in maroon. Non-significant values are left blank and represented in white. The gene list is based on available literature. **C.** Heat map representing protein level fold change of Golgi apparatus structural and functional proteins from mass spectrometry measurement of total cell extracts from Fig S3C prepared from RPE1 cells treated with siFBL or siNON controls for 48- and 96 hours. The scale bar ranges from 0 to 2-fold change and proteins follow a p-value cutoff of 0.05. Out of scale values are represented in maroon. The protein list is based on available literature. **D.** Immunoblots of total cell lysates prepared from RPE1 cells treated with the indicated siRNAs for 96 hours (siNON, siFBL, siUBF, siUTP18) and probed with antibodies against GM130. GAPDH controls equal loading. **E.** Representative confocal microscopic images of RPE1 cells treated with the indicated siRNAs for 96 hours (siNON, siFBL, siUBF, siUTP18), and co-stained for the cis-Golgi marker protein GM130 (green) and DAPI (blue). Images are representative of N=4 biological replicates. Scale bar: 10 µm. **F.** Bar graph showing data from Fig 4E, measuring the percentage (%) of RPE1 cells classified based on GM130 stainings with an enlarged Golgi using manual annotation, following treatment with the indicated siRNA for 96 hours (siNON, siFBL, siUBF, siUTP18). Mean with SD is plotted from N=4 biological replicates. Statistical analysis was performed by an ordinary one-way Anova test with Tukey’s multiple comparison. Asterisks denote p value (****, <0.0001). **G.** Heat map representing genes fold change of endosome and lysosome structural and functional components from RNA-seq datasets of RPE1 cells treated with siFBL, siUTP18 and siNON controls for 48- and 96 hours. The scale bar ranges from 0 to 2-fold change and genes follow a p-value cutoff of 0.05. Out of scale values are represented in maroon. Non-significant values are left blank and represented in white. The gene list is based on available literature. **H.** Heat map representing protein level fold change of endosome and lysosome structural and functional proteins measured by mass spectrometry of extract prepared from RPE1 cells treated with siFBL or siNON controls for 48- or 96 hours. The scale bar ranges from 0 to 2-fold change and proteins follow a p-value cutoff of 0.05. Out of scale values are represented in maroon. The protein list is based on available literature. **I.** Representative confocal microscopic images of RPE1 cells treated with siNON and siFBL for 96 hours, and co-stained for the endosomal marker protein EEA1 (green) and DAPI (blue). Images are representative of N=2 biological replicates. Scale bar: 10 µm. **J.** Representative confocal microscopic images of RPE1 cells treated with siNON and siFBL for 96 hours, and co-stained for the lysosomal marker protein TFEB (green) and DAPI (blue). Images are representative of N=3 biological replicates. Scale bar: 10 µm. **K.** Representative confocal microscopic images of RPE1 cells treated with siNON and siFBL for 96 hours, and co-stained for the endosomal and lysosomal marker protein LAMP1 (green) and DAPI (blue). Images are representative of N=3 biological replicates. Scale bar: 10 µm. **L.** Bar graph showing the percentage (%) of cells classified with one nucleolus phenotype (counted as 1 or 2 nucleoli) with small (comparable to siNON condition) Golgi, many nucleoli (>2 nucleoli) with small Golgi, one nucleolus with big (enlarged) Golgi and many nucleoli with big Golgi using manual annotation following treatment with indicated siRNA for 48-, 96- and 192 hours (siNON, siFBL, siUBF, siUTP18), as observed with NPM and GM130 staining. Mean with SEM is plotted from N=3 biological replicates. **M.** Bar graph showing the percentage (%) of cells manually classified with one nucleolus phenotype (counted as 1 or 2 nucleoli) with small (comparable to siNON conditions) endosomes, many nucleoli (>2 nucleoli) with small endosomes, one nucleolus with big (enlarged and many) endosomes and many nucleoli with big endosomes in RPE1 cells treated with the indicated siRNA for 48-, 96- and 192 hours (siNON, siFBL, siUBF, siUTP18), as observed with NPM and EEA1 staining. The mean with SEM is plotted from N=3 biological replicates.

### 5S RNP-mediated TP53 activation triggers profound nucleolar and cytoplasmic changes

To dissect the molecular mechanisms responsible for TP53 activation upon FBL depletion, we performed co-depletion experiments of FBL and major ribosomal proteins involved in TP53 activation and assessed rescue of the cellular phenotypes associated with nucleolar defects. Along with other ribosomal proteins, RPL5 and RPL11 are primarily known to activate TP53 upon ribotoxic stress by binding its negative regulators MDM2 and MDM4^42,43,55^. As expected, co-depletion of RPL5 and RPL11 along with FBL abolished TP53 stabilization and CDKN1A upregulation **(Fig 5A and S5A-B)**. Interestingly, Ribosome Production Factor 2 (RPF2), an essential protein in the 5S RNP complex responsible for RPL5-RPL11 incorporation into pre-60S particles^56,57^, was upregulated at the earliest stress time point **(Fig 3C)**. Although the role of RPF2 in regulating TP53 signaling upon ribotoxic stress remains elusive, cells lacking RPF2 exhibited TP53 stabilization and CDKN1A upregulation **(Fig 5B and S5C)**. These findings demonstrate that 5S RNP defects and impaired incorporation of RPL5-RPL11 into pre-ribosomal particle initiates TP53 stabilization and subsequent CDKN1A expression.

**Figure 5:**
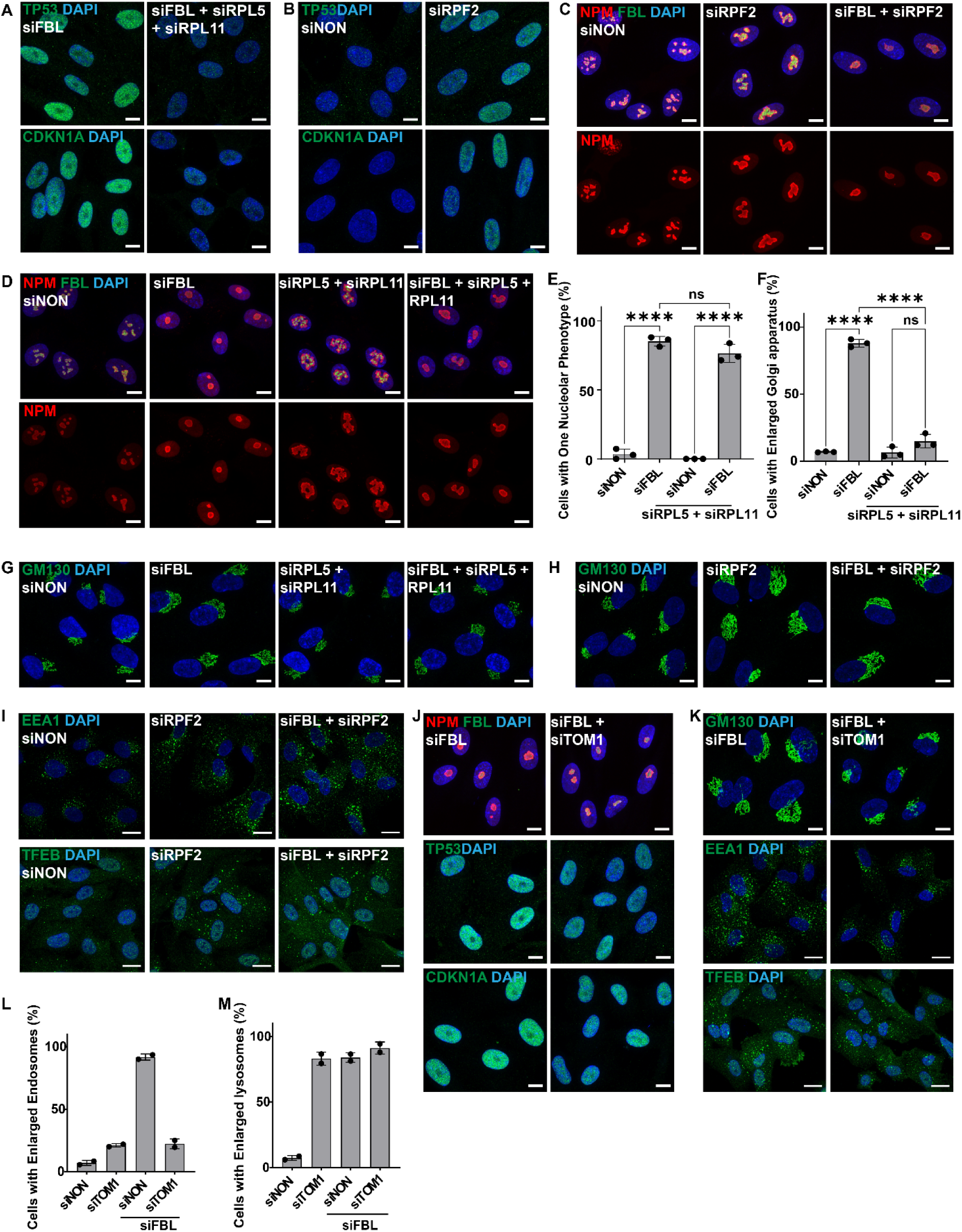
Cytoplasmic rearrangements upon chronic nucleolar defects depend on 5S RNP mediated TP53 activation. **A.** Representative confocal microscopic images showing RPE1 cells treated with the indicated single or double siRNAs for 96 hours (siFBL, siFBL+ siRPL5 + siRPL11). Cells in the top panel are co-stained for TP53 (green) and DAPI (blue) and cells in the bottom panel for CDKN1A (green) and DAPI (blue). Images are representative of N=2 biological replicates. Scale bar: 10 µm. **B.** Representative confocal microscopic images showing RPE1 cells treated as indicated with single or double siRNAs for 96 hours (siNON, siRPF2). Cells in the top panel are co-stained for TP53 (green) and DAPI (blue) and cells in bottom panel for CDKN1A (green) and DAPI (blue). Images are representative of N=2 biological replicates. Scale bar: 10 µm. **C.** Representative confocal microscopic images showing RPE1 cells following treatment with the indicated siRNA for 96 hours (siNON, siRPF2, siFBL + siRPF2), and co-stained for NPM (red), FBL (green) and DAPI (blue). The top row displays merged co-localization and the bottom row NPM localization. Images are representative of N=2 biological replicates. Scale bar: 10 µm. **D.** Representative confocal microscopic images showing RPE1 cells treated with the indicated siRNA for 96 hours (siNON, siFBL, siRPL5 + siRPL11, siFBL+ siRPL5 + siRPL11) and co-stained for NPM (red), FBL (green) and DAPI (blue). Cells in the top row display merged co-localization and cells in the bottom row NPM localization. Images are representative of N=3 biological replicates. Scale bar: 10 µm. **E.** Bar graph showing data from Fig 5D, measuring the percentage (%) of RPE1 cells classified as having 1 spherical nucleolus (indicated as one nucleolar phenotype) at 96 hours of the indicated siRNA treatments, as observed with NPM staining. The mean with SD is plotted from N=3 biological replicates. Statistical analysis was performed by a one-way Anova test with Dunnett’s multiple comparison. Asterisks denote p value (****, <0.0001). ns, non-significant. **F.** Bar graph showing data from Fig 5G, measuring the percentage (%) of RPE1 cells classified as having an enlarged Golgi at 96 hours siRNA treatment, as observed with GM130 staining. The mean with SD is plotted from N=3 biological replicates. Statistical analysis was performed by a one-way Anova test with Dunnett’s multiple comparison. Asterisks denote p value (****, <0.0001). ns, non-significant. **G.** Representative confocal microscopic images showing RPE1 cells treated with the indicated siRNA for 96 hours (siNON, siFBL, siRPL5 + siRPL11, siFBL+ siRPL5 + siRPL11) and co-stained with the cis-Golgi marker GM130 (green) and DAPI (blue). Images are representative of N=3 biological replicates. Scale bar: 10 µm. **H.** Representative confocal microscopic images showing RPE1 cells treated with the indicated siRNA for 96 hours (siNON, siRPF2, siFBL + siRPF2) and co-stained with the cis-Golgi marker GM130 (green) and DAPI (blue). Images are representative of N=2 biological replicates. Scale bar: 10 µm. **I.** Representative confocal microscopic images showing RPE1 cells treated with the indicated siRNA for 96 hours (siNON, siRPF2, siFBL + siRPF2) and co-stained for endosomal marker EEA1 (green) and DAPI (blue) (top panel) or the lysosomal marker TFEB (green) and DAPI (blue) (bottom panel). Images are representative of N=2 biological replicates. Scale bar: 20 µm. **J.** Representative confocal microscopic images showing RPE1 cells treated with the indicated siRNA for 96 hours (siFBL, siFBL + siTOM1) and co-stained for NPM (red), FBL (green) and DAPI (blue) (top panel), or TP53 (green) and DAPI (blue) (middle panel) or CDKN1A (green) and DAPI (blue) (bottom panel). Images are representative of N=2 biological replicates. Scale bar: 20 µm. **K.** Representative confocal microscopic images showing RPE1 cells treated with the indicated siRNA for 96 hours (siFBL, siFBL + siTOM1) and co-stained for the cis-Golgi marker GM130 (green) and DAPI (blue) (top panel), for the endosomal marker EEA1 (green) and DAPI (blue) (middle panel) and the lysosomal marker TFEB (green) and DAPI (blue) (bottom panel). Images are representative of N=2 biological replicates. Scale bars: 10 µm, 20 µm, 20 µm. **L.** Bar graph showing data from Fig 5K measuring the percentage (%) of RPE1 cells classified as having bigger and an increased number of endosomes (indicated as enlarged endosomal phenotype) at 96 hours siRNA treatment, as observed with EEA1 staining. The mean with SD is plotted from N=2 biological replicates. **M.** Bar graph showing data from Fig 5K measuring the percentage (%) of RPE1 cells classified as having bigger and an increased number of lysosomes (indicated as enlarged lysosomal phenotype) at 96 hours siRNA treatment, as observed with TFEB staining. The mean with SD is plotted from N=2 biological replicates.

Interestingly, depletion of all three 5S RNP components led to an aberrant nucleolar morphology characterized by size increase, irregular shape and GC fusion and enlargement **(Fig 5C-E and S5D-F),** which is distinct from the FBL-depletion phenotype. Surprisingly, unlike TP53, RPL5 and RPL11 co-depletion failed to rescue the one-nucleolar morphology **(S3I-K)**, implying that additional TP53-depending signaling pathways regulate nucleolar morphology upon loss of nucleolar proteins. In addition, their co-depletion with FBL resulted in a nucleolar morphology indistinguishable from FBL alone **(Fig 5C-D and S5D)**, further validating that nucleolar defects arising from FBL depletion are upstream of the 5S RNP complex in nucleolar ribosome biogenesis. Indeed, these nucleolar defects likely arise at different steps of rRNA processing and pre-ribosomal assembly^16^.

Interestingly, co-depletion of RPL5 and RPL11 along with FBL abolished cytoplasmic rearrangements such as Golgi enlargement **(Fig 5F-G and S5G)**, indicating that these responses require robust TP53 stabilization. To our surprise, RPF2 depletion alone already caused cytoplasmic rearrangements similar to the FBL-like enlarged Golgi and bigger endo-lysosomes **(Fig 5H-I)**. Co-depletion of RPF2 and FBL failed to rescue the cytoplasmic phenotypes, confirming that both proteins are part of a common signaling axis. Together these results demonstrate that the cytoplasmic rearrangements originate from 5S RNP defects and subsequent TP53 activation. The uncoupling of nucleolar morphology defects from cytoplasmic organelle rewiring suggests that the nucleolar morphology changes arise from ribosome biogenesis defects in the nucleolus, which occur upstream of TP53 activation, whereas the cytoplasmic rearrangements are subsequent downstream consequences of TP53 activation.

Another possible TP53 dependent regulator of the cytoplasmic rearrangements is Target of Myb1 membrane trafficking protein 1 (TOM1), a reported endosomal protein known to play a crucial role in vesicular trafficking^58,59^. TOM1 showed persistent upregulation in the proteomic datasets **(Fig 4H)**, possibly through its putative upstream regulators MYBBP1A and MYB, which are involved in TP53 activation^60^. As expected, TOM1 depletion alone or combined with FBL depletion did not alter nucleolar and nuclear phenotypes **(Fig 5J and S5H)**. However, cells lacking TOM1 showed a striking increase in the number of TFEB-stained lysosomes but not EEA1-stained endosomes **(Fig S5H)**, and co-depletion of TOM1 and FBL abrogated the Golgi and endosome defects but not the lysosome phenotype **(Fig 5K-M).** Hence, TOM1 functionally links nucleolar defects to changes in cytoplasmic organelles. Additionally, CCN2 was persistently upregulated in our datasets, and its depletion resulted in increased cell size, but no other phenotypes were detected even when co-depleted with FBL **(Fig S5I)**. Taken together, these results suggest crosstalk among TP53 pathways, in particular nucleolar defects and cytoplasmic regulators of organellar homeostasis.

### Golgi and vesicle transport pathways are causally linked to prolonged nucleolar defects

To determine whether the cytoplasmic alterations are functionally relevant for cells adapting to nucleolar defects, we performed an unbiased genome-wide CRISPRi screen, searching for genes that suppress or enhance proliferation of FBL-depleted cells. The engineered dCAS9-KRAB RPE1 cells exhibit proliferation and nucleolar defects comparable to those of wild type RPE1 cells **(Fig S6A-C)**. We used the human genome-wide CRISPRi-v2 lentiviral library consisting of 102’640 gRNAs that inhibit gene transcription as well as 1’895 non-targeting controls^61^. Following lentiviral transduction, dCAS9-KRAB RPE1 cells were puromycin-selected and cultured for 10 days to ensure robust knockdowns. Fibrillarin was then depleted by siRNA, and differences in sgRNA abundance between FBL-depleted and control (siNON) cells were quantified by deep sequencing at 120h, 240h and 360h post siRNA treatment **(Fig 6A)**. For validation, we first examined the 1’750 genes with the lowest sequencing read counts across all siNON-treated conditions comprising approximately 10% of the library, and found that more than two-thirds, 1’176 genes, were common among all 3 time points **(Fig S6D)**. Gene enrichment analysis confirmed loss of essential genes involved in DNA replication, ribosome biogenesis, cell cycle and splicing processes **(Fig S6E-F)**. Plotting the relative log2-fold change normalized to the condition mean distribution of sequencing reads revealed condition- and time point specific effects of all 17’592 detected genes in cell survival and proliferation **(Fig 6B)**. Grouping the 2’000 genes with the highest standard deviation among all six experimental conditions encompassing two knockdowns and three time points led to six functional clusters identifying genes enhancing or suppressing proliferation and/or cell survival caused by FBL depletion **(Fig 6C)**. Clusters 1 and 6 were genes that altered proliferation independently of FBL and thus not further investigated. Of note, clusters 3 and 4 comprise genes synthetically lethal with FBL depletion at later time points, including 240h and 360h **(Fig 6D)**. Gene enrichment analysis showed over-representation of genes involved in rRNA processing, ribosome biogenesis and cytoplasmic ribosomes, validating the robustness and biological relevance of the CRISPRi screen **(Fig 6D)**. Interestingly, cluster 2 and 5 encompass genes that increase survival and/or proliferation of cells with nucleolar defects and comprise a class of genes with roles in overall nucleolar stress response and cell cycle progression **(Fig 6E)**. To validate the functional relevance of these candidates, we performed systematic co-depletion of FBL with promising hits from the CRISPRi screen and assessed their possible roles in stress signaling from the nucleolus to Golgi and endosomal trafficking. As expected, TP53 and CDKN1A were among the top hits, improving proliferation of FBL-depleted cells throughout the time course, and already detectable at the 120h time point (**Fig 6B**). Surprisingly, co-depletion of FBL and CDKN1A not only restored cell proliferation but also abrogated the one-nucleolus and enlarged Golgi phenotypes without altering TP53 stabilization **(Fig S6G-J)**, implying that nucleolar morphology and the cytoplasmic changes can be indirectly abolished by cell cycle rescue.

**Figure 6:**
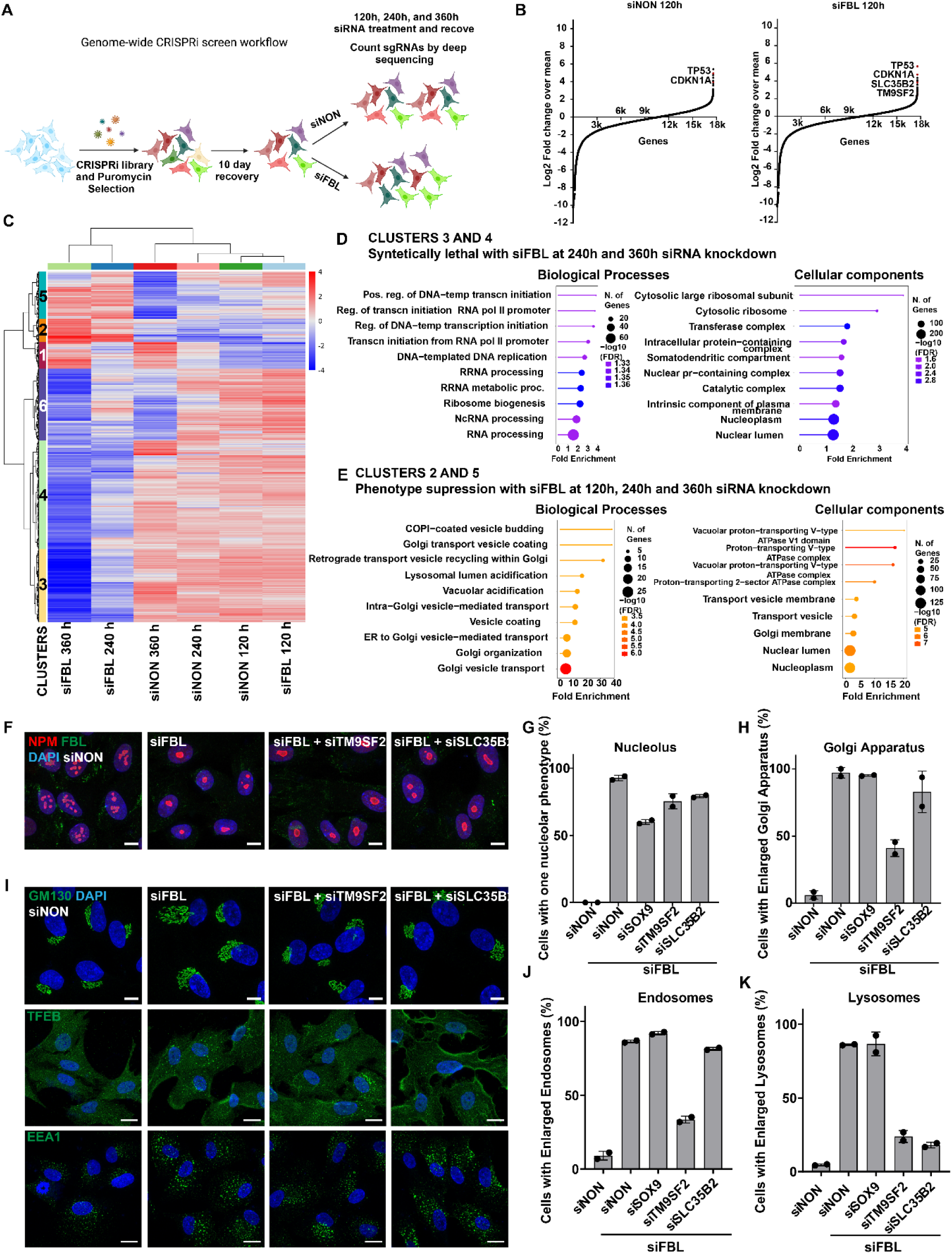
Genome-wide CRISPRi screen identifies functional interactors of chronic nucleolar defects. **A.** Schematic showing the experimental setup of the genome-wide CRISPRi screen. A human genome-wide lentiviral library consisting of >100,000 gRNAs that inhibit transcription of the selected gene was used. RPE1 cells were cultured in selection media for 10 days prior to treatment with siNON controls or siFBL. sgRNA abundance was quantified by deep sequencing. **B.** Map showing rank ordering all 17592 genes, identified by deep sequencing of CRISPRi-treated FBL-depleted cells. Plot represents log2 converted values representing the fold change comparing the indicated condition and time-point specific column mean for siFBL and siNON control cells at 120 hours of siRNA treatment. **C.** Heatmap representing 2000 genes with high standard deviation among six experimental conditions identified by deep sequencing. The threshold for std. dev. of the log_2_ signal across samples is 1.13. The heatmap is classified into six functional clusters based on condition dependent differences in proliferation and/or survival resulting from the detected sgRNA abundance. The differences are plotted on a scale of -4 to +4. **D.** Lollipop chart of biological processes (left panel) and cellular component (right panel) enrichment analysis using ShinyGO from the CRISPRi dataset selecting genes belonging to cluster 3 and 4 from the heatmap in Fig 6C. Pathways over fold enrichment were plotted with -log10FDR using the indicated color scheme. Silencing of these genes results in reduced proliferation/ survival when combined with siFBL and are thus referred to as synthetically lethal. Number of genes: 1035 and FDR cutoff: 0.01. **E.** Lollipop chart of biological processes (left panel) and cellular component (right panel) enrichment analysis using ShinyGO from the CRISPRi dataset selecting genes belonging to cluster 2 and 5 from the heatmap in Fig 6C. Pathways over fold enrichment were plotted with -log10FDR using the indicated color scheme. Silencing of these genes results in increased proliferation/ survival when combined with siFBL and are thus referred to as suppressors. Number of genes: 399 and FDR cutoff: 0.01. **F.** Representative confocal microscopic images of RPE1 cells following treatment with indicated siRNAs for 96 hours (siNON, siFBL, siTM9SF2 and siSLC35B2), and co-stained for NPM (red), FBL (green) and DAPI (blue). Images are representative of two biological replicates (N=2). Scale bar: 10 µm. **G.** Bar graph showing data from Fig 6F and S6M measuring the percentage (%) of cells classified with one nucleolar phenotype following treatment with the indicated siRNA for 96 hours (siNON, siFBL, siSOX9, siTM9SF2 and siSLC35B2), as observed with NPM staining. Mean with SD is plotted from N=2 biological replicates. **H.** Bar graph showing data from Fig 6I and S6M measuring the percentage (%) of cells classified with the enlarged Golgi phenotype of RPE1 cells treated with the indicated siRNA for 96 hours (siNON, siFBL, siSOX9, siTM9SF2 and siSLC35B2), as observed with GM130 staining. Note that co-depletion of TM9SF2 and FBL shows rescue of the enlarged Golgi phenotype. The mean with SD is plotted from N=2 biological replicates. **I.** Representative confocal microscopic images of RPE1 cells treated with the indicated siRNAs for 96 hours (siNON, siFBL, siTM9SF2 and siSLC35B2). The top row shows co-staining of the cis-Golgi protein GM130 (green) and DAPI (blue), the middle row co-staining of the lysosomal protein TFEB (green) and DAPI (blue), and the bottom row merged co-staining of the endosomal protein EEA1 (green) and DAPI (blue). Images are representative of N=2 biological replicates. Scale bars: 10 µm, 20 µm and 20 µm. **J.** Bar graph showing data from Fig 6I and S6M measuring the percentage (%) of cells classified with the enlarged endosomal phenotype of RPE1 cells treated with the indicated siRNA for 96 hours (siNON, siFBL, siSOX9, siTM9SF2 and siSLC35B2), as observed with EEA1 staining. Note that co-depletion of TM9SF2 and FBL show rescue of the enlarged endosomal phenotype. The mean with SD is plotted from N=2 biological replicates. **K.** Bar graph showing data from Fig 6I and S6M measuring the percentage (%) of cells classified with the enlarged lysosomal phenotype of RPE1 cells treated with the indicated siRNA for 96 hours (siNON, siFBL, siSOX9, siTM9SF2 and siSLC35B2), as observed with TFEB staining. Note that co-depletion of TM9SF2 or SLC35B2 and FBL show rescue of the enlarged lysosomal phenotype. The mean with SD is plotted from N=2 biological replicates.

Importantly, the gene enrichment analysis showed that cluster 2 and 5 contain genes involved in Golgi organization, Golgi vesicle transport, lysosomal and vacuolar acidification **(Fig 6E)**, including the uncharacterized Golgi and endosome localized protein TM9SF2^62,63^ and the Golgi membrane transporter SLC35B2^64^. Knockdown of these two genes did not significantly alter the nucleolar phenotypes and TP53 stabilization of FBL-depleted cells **(Fig 6F-G and S6K)**. However, when co-depleted with FBL, TM9SF2 rescued the enlarged Golgi structures and endo-lysosomal phenotypes whereas SLC35B2 only rescued lysosomal phenotypes **(Fig 6H-K and S6L)**. Interestingly, the transcription factor SOX9, involved in expressing a putative ECM gene signature, was upregulated in our datasets but didn’t rescue proliferation at early and mid-time points in the CRISPRi screen. Instead, SOX9 depletion alone led to an enlarged cell size **(Fig S6M)**, indicative of senescence signaling, while co-depletion of SOX9 with FBL showed no rescue of the cellular phenotypes **(Fig S6M)**. As SLC35B2 and TM9SF2 were identified in the functional CRISPR screen, we speculate that an enlarged Golgi apparatus and increased endo-lysosomes are biologically important for cell cycle arrest and to attain putative cellular quiescence. Taken together, we conclude that these cytoplasmic components respond to prolonged nucleolar defects downstream of TP53 without affecting nucleolar morphology, implying that Golgi expansion and lysosomal vesicular transport constitute key pathway in attaining or maintaining cellular quiescence as a response to persistent loss of ribosome biogenesis and nucleolar function.

### Chronic nucleolar stress upregulates secretion of ECM scaffolding and regulatory proteins

In contrast to early mediators of the nucleolar stress response, several secretory proteins and ECM components like TGFB2, CCN2, CPA4, TOM1 and COL4A2 showed persistent upregulation after FBL depletion. Moreover, upregulation of collagen genes remained detectable even beyond the recovery time point of 192h **(Fig 7A).** This analysis suggests that nucleolar defects initiate a coordinated gene expression program, culminating in the upregulation and secretion of collagen and other extracellular matrix components to allow cellular adaptation. Consistent with this notion, TEM images of FBL-depleted cells uncovered a striking increase in secretory vesicles **(Fig 7B)**, and immunofluorescence and confocal imaging confirmed increased production of collagen (COL4A1) **(Fig 7C)**. We also stained for fibronectin (FN1), a prominent ECM component with distinguishable intra- and extracellular forms. While intracellular FN1 marks Golgi and cytoplasm, extracellular FN1 can be distinguished as sharp fiber-like structures. Both intra and extracellular FN1 showed prominent upregulation upon FBL depletion **(Fig 7D)**, and COL4A1 and FN1 were upregulated, at least in part, by a TP53-dependent mechanism **(Fig 7C-D and Fig S7A-B)**. To directly validate increased secretion, we used mass spectrometry to analyze cell culture supernatants. We first measured the secretome with FCS present in the media at 48h and 96h post FBL knockdown, and identified many upregulated components such as CCN2, FN1 and FBN1 **(Fig S7C)**. Gene ontology analysis upon FBL depletion revealed enriched genes involved in wound healing, angiogenesis and cell adhesion belonging to collagen containing matrix, extracellular matrix and vesicles, validating robustness of the assay and confirming secretory adaptations upon chronic nucleolar defects **(Fig S7D)**. As we were unable to distinguish bovine and human TGFB2 in the secretome, we repeated the measurements after removing FCS from the media 24h prior to harvesting. While control (siNON) cells became more secretory upon FCS removal^65^, significant upregulation of CCN2, COL4A1-2, CPA4 and TGFB2 was detected in FBL-depleted cells **(Fig 7E)**. CCN2 functions as an accessory regulator of the TGFB2 pathway, but its depletion did not rescue the stress phenotypes associated with FBL depletion **(Fig S6M)**. Interestingly, even though TP53 activation via the RPL5-RPL11 pathway drives cellular senescence^46^, we did not observe a canonical senescence-associated secretion profile in FBL-depleted cells. Instead, the secretome of FBL-depleted cells mainly comprised components regulating ECM remodeling and TGF-B signaling, in addition to a few senescence-associated signatures. Transcriptionally, these cells upregulated the expression of pro-EMT cytokines like IL6 and IL33, whereas other cytokines remained poorly expressed. Cells also upregulated the expression of many IL receptors without changing the levels of the corresponding ligands, indicating a quiescence or poised state **(Fig S7E)**. Taken together, we conclude that chronic nucleolar dysfunction reorganizes cellular secretion leading to changes in cytoplasmic and extracellular signaling resembling cellular quiescence.

**Figure 7:**
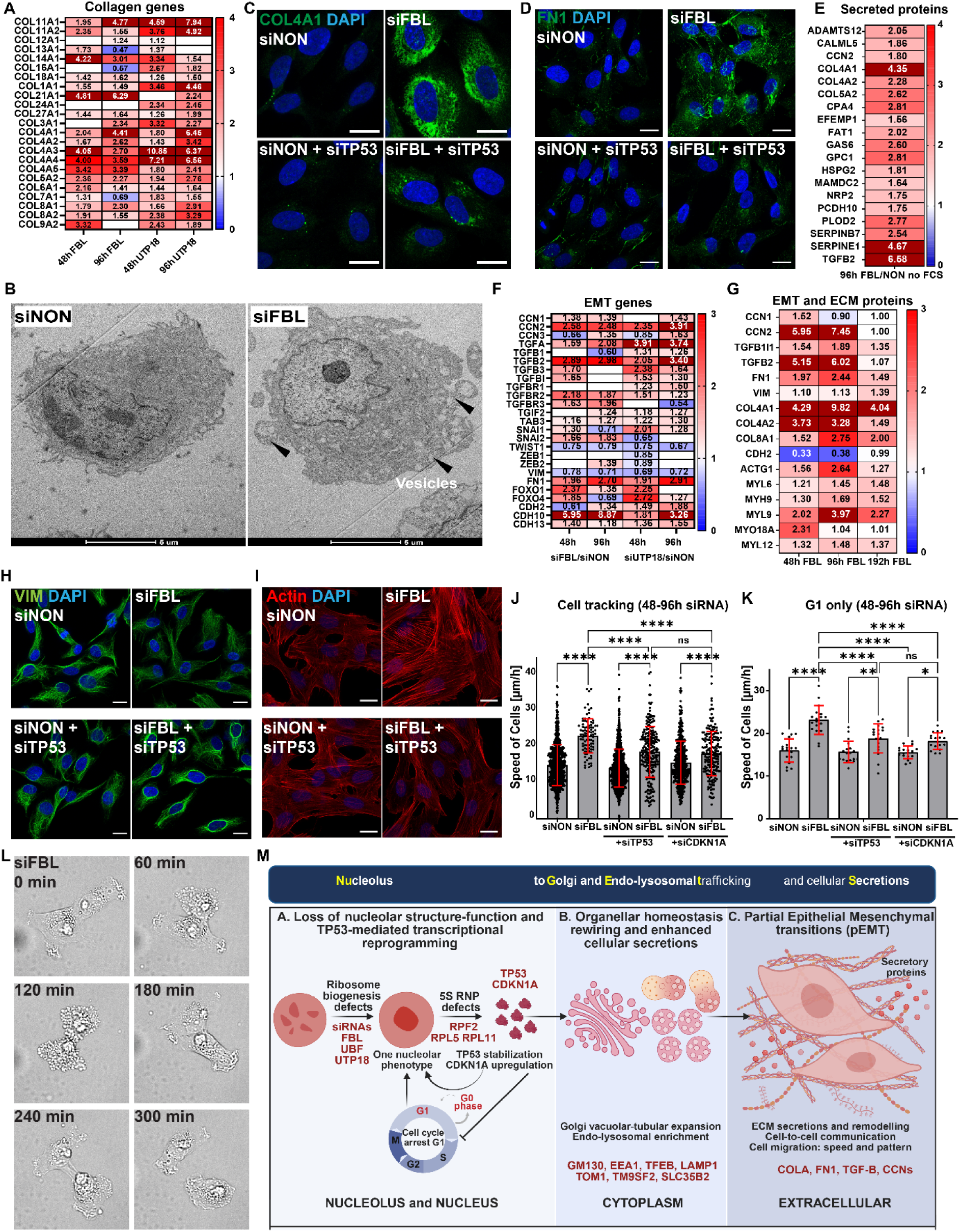
Chronic nucleolar defects lead to cellular quiescence and EMT like phenotypes. **A.** Heat map representing genes fold change of different classes of collagen proteins from RNA-seq datasets of RPE1 cells treated with siFBL, siUTP18 or siNON controls for 48- and 96 hours. The scale bar ranges from 0 to 4-fold change and genes follow a p-value cutoff of 0.05. Out of scale values are represented in maroon. Non-significant values are left blank and represented in white. The gene list is based on available literature. **B.** Transmission electron micrographs showing cellular architecture and cytoplasmic features of RPE1 cells treated with siFBL and siNON controls for 96 hours. Arrows mark accumulation and exocytosis of cytoplasmic vesicles. Scale bar: 5 µm. **C.** Representative confocal microscopic images of RPE1 cells treated with the indicated single or double siRNAs for 96 hours (siNON, siFBL, siNON+siTP53, siFBL+siTP53), and co-stained for collagen COL4A1 (green) and DAPI (blue). Images are representative of N=4 biological replicates. Scale bar: 20 µm. **D.** Representative confocal microscopic images of RPE1 cells treated with the indicated single or double siRNAs for 96 hours (siNON, siFBL, siNON+siTP53, siFBL+siTP53), and co-stained for fibronectin FN1 (green) and DAPI (blue). Images are representative of N=3 biological replicates. Scale bar: 20 µm. **E.** Heat map representing protein level fold change of prominent secreted proteins identified by mass spectrometry from cell culture supernatants (secretome) obtained from RPE1 cells treated with siFBL or siNON controls for 96 hours. The cells were cultured 24 hours without fetal calf serum (FCS) prior to collection. The scale bar ranges from 0 to 4-fold change and proteins follow a p-value cutoff of 0.05. Out of scale values are represented in maroon. **F.** Heat map representing genes fold change of Epithelial to Mesenchymal Transition (EMT) associated proteins from RNA-seq dataset of RPE1 cells treated with siFBL, siUTP18 or siNON controls for 48- and 96 hours. The scale bar ranges from 0 to 3-fold change and genes follow a p-value cutoff of 0.05. Out of scale values are represented in maroon. Non-significant values are left blank and represented in white. The gene list is based on available literature. **G.** Heat map representing protein level fold change of Extracellular matrix (ECM) and Epithelial to Mesenchymal Transition (EMT) associated proteins identified by mass spectrometry in extracts prepared from RPE1 cells treated with siFBL or siNON controls for 48-, 96- and 192 hours. The scale bar ranges from 0 to 3-fold change and proteins follow a p-value cutoff of 0.05. Out of scale values are represented in maroon. The protein list is based on available literature. **H.** Representative confocal microscopic images of RPE1 cells treated with the indicated single or double siRNAs for 96 hours (siNON, siFBL, siNON+siTP53, siFBL+siTP53), and co-stained for the intermediate filament protein Vimentin (VIM, green) and DAPI (blue). Images are representative of N=4 biological replicates. Scale bar: 20 µm. **I.** Representative confocal microscopic images of RPE1 cells treated with the indicated single or double siRNAs for 96 hours (siNON, siFBL, siNON+siTP53, siFBL+siTP53), and co-stained for actin stress fibers using phalloidin (red) and DAPI (blue). Images are representative of N=4 biological replicates. Scale bar: 20 µm. **J.** Bar graph showing the migration speed derived from live-cell imaging of FUCCI-RPE1 cells treated with the indicated siRNA (siNON, siFBL, siTP53, siCDKN1A, siFBL + siTP53, siFBL + siCDKN1A) for 48 to 96 hours. 30 cells per condition and replicate are chosen at the 48-hour time point and used together with their progeny for continuous tracking of cell mobility. The mean with SD is plotted for N=3 biological replicates. Statistical analysis was performed by an Ordinary one-way Anova test with Tukey’s multiple comparison. Asterisks denote p value (****, <0.0001; **, <0.01; *, <0.05). ns, non-significant. **K.** Bar graph showing of the speed of migrating cells derived from live-cell imaging of FUCCI-RPE1 cells in the G0/G1 cell-cycle phase, treated with the indicated siRNA (siNON, siFBL, siTP53, siCDKN1A, siFBL + siTP53, siFBL + siCDKN1A) for 48 hours to 96 hour time points. Each data point represents the average speed of five starting cells and their progeny imaged in single imaging frames. The mean with SD is plotted for N=3 biological replicates. Statistical analysis was performed by an Ordinary one-way Anova test with Tukey’s multiple comparison. Asterisks denote p value (****, <0.0001). ns, non-significant. **L.** Brightfield snapshots derived from live-cell imaging of FUCCI-RPE1 cells treated with siFBL showing secretory and cell-cell communication behavior. Images were taken every 60 minutes and are presented over a period of 300 minutes starting from observed cell-cell contacts. **M.** Schematic illustration summarizing the major findings. Nucleolar 5S RNP assembly defects caused by non-lethal nucleolar changes trigger TP53 activation, which causes cell cycle arrest, and rewires cytoplasmic organelles and promotes secretion and ECM remodeling. In turn, TP53 activity also alters nucleolar morphology. This signaling network termed NuGETS allows cells to adopt EMT-like behavior, characteristic of ribosomopathies.

### Chronic nucleolar defects alter cell migration and cell-cell communication patterns

To better understand the function of these ECM and secretome changes, we searched for gene regulatory networks linking cellular secretion to cell proliferation and stress adaptation. Interestingly, we detected significant upregulation of YAP/TAZ-TEAD signaling and negative regulators of WNT in cells lacking FBL, indicative of a mechanosensing pathway coordinating cellular quiescence and secretion **(Fig S7F-G)**. Moreover, we found that many genes involved in the epithelial-to-mesenchymal transition (EMT) such as TGFB2, TGFBR, CCN2 and cadherins were significantly upregulated at the transcriptome and proteome levels in cells lacking FBL or UTP18 **(Fig 7F-G)**. We also identified that the cancer-associated EMT transcription factors SNAIL1^66^ and SNAIL2^67^ were upregulated upon prolonged nucleolar defects but not the canonical EMT-TFs regulating developmental programs such as TWIST1 and ZEB1 **(Fig 7F)**. To determine whether cells with nucleolar defects develop an EMT-like phenotype, we stained the intermediate filament vimentin (VIM) and tubulin and observed an altered squamous-like cell shape **(Fig 7H and S7H-I)**. Similarly, cells lacking FBL or UTP18 showed an increase in the number and intensity of actin stress fibers **(Fig 7I and S7I-J)**, in contrast to UBF depletion **(Fig S7G)**. These cytoskeletal and cell shape changes were abrogated if TP53 was co-depleted **(Fig 7H-I and Fig S7H-J)**. Tracking individual RPE1-FUCCI cells by live cell imaging revealed that these cytoskeletal rearrangements alter cell migration **(Fig 3G-H and Fig S3H)**, and the migration speed prominently increased after 48 to 96h of siRNA knockdown **(Fig 7J and Fig S7K)**. These cell migration changes were partially dependent on TP53 and its downstream target CDKN1A **(Fig 7J-K and Fig S7K-L)**, although increased migration speed was also observed when analyzing only G1 cells **(Fig 7K and S7L)**. Surprisingly, live cell imaging of FBL-depleted cells also revealed striking changes in cell-cell interactions. The RPE1 cells showed an oscillating secretory phenotype that was synchronized with and dependent on neighboring cells **(Fig 7L)**. Overall, these results demonstrate that a nucleolar stress response pathway promotes activation of Golgi and trafficking organelles, rewiring secretion and rearranging the ECM to achieve altered cell migration and cell-cell communication patterns, resembling the EMT process.

## Discussion

Using a multi-omics approach, we uncovered an adaptive stress-response network termed NuGETS (<u>Nu</u>cleolus to <u>G</u>olgi, <u>E</u>ndo-lysosomal <u>T</u>rafficking and <u>S</u>ecretion), originating from persistent nucleolar dysfunction **(Fig 7M)**. Cells lacking nucleolar components undergo a TP53-dependent de-differentiation program resembling partial EMT (pEMT) and cellular quiescence. Consequently, cells show an enlarged Golgi apparatus and upregulate endo-lysosomal vesicles, leading to increased secretion of ECM components and signaling molecules. Together, these responses promote cell survival and a pro-metastatic behavior with increased cell migration, characteristic of cancer types caused by ribosomopathies.

### Distinct cellular responses upon acute and prolonged nucleolar stress conditions

The notion that ribosome biogenesis defects lead to nucleolar shut-off of rRNA transcription and processing, inhibition of Pol II transcription in the nucleus and cytoplasmic translation shut-down^34,38^ is primarily derived from data with acute stressors such as heat-shock^68^ or chemicals including the Pol I inhibitor ActD, the metabolic poison arsenate or the DNA-damaging drug CPT^33,34^. These acute conditions activate general stress pathways and induce secondary effects, which can trigger cellular senescence and apoptosis^46,69–71^. While these studies revealed important insight into rapid mechanisms protecting cell integrity, prolonged stress conditions such as nutrient starvation allow cells to adapt and maintain sufficient cytoplasmic translation capacity to carry out essential functions^72,73^. However, nutrient deprivation substantially alters cell physiology, making it difficult to disentangle direct and indirect effects. Similarly, experiments performed by depleting essential ribosomal proteins disrupt cytoplasmic translation and accumulate free ribosomal proteins and/or subcomplexes^74,75^, which result in cellular apoptosis or senescence^46,75^. To circumvent these complications, we investigated the cellular response to the gradual loss of key nucleolar proteins giving rise to specific structural and functional nucleolar defects. Interestingly, these experiments reveal that cells have a large buffering capacity to preserve sufficient Pol II transcription and ribosome biogenesis to sustain cytoplasmic translation, consistent with previous reports^48,76^. Moreover, cells can counteract dilution of cytoplasmic ribosomes by arresting cell division and possibly inhibiting ribosome degradation^73^. Together, these mechanisms compensate for reduced ribosome biosynthesis and ensure cell survival to allow adaptation. As a results cells exposed to prolonged but mild nucleolar defects adapt and show a non-canonical senescence signature including reversible cell cycle arrest and cell enlargement.

### Unexpected crosstalk among nucleolar proteins

The metazoan nucleolus functions as a membraneless organelle and confines more than 600 proteins that cooperate to orchestrate ribosome biogenesis^77,78^. However, the different nucleolar constituents do not work independently but rather influence each other to maintain homeostasis^49^. For instance, depletion of FBL or UTP18 results in transcriptional downregulation and reduced protein levels of several key nucleolar proteins such as nucleolin, UTP complexes and nucleophosmin. Similarly, the levels and localization of nucleolar resident proteins is significantly altered in response to nucleolar defects induced by depletion of seemingly different structural and functional nucleolar components^18^. Thus, caution is warranted if the function of specific nucleolar constituents is assigned solely based on knock-out phenotypes. Such unexpected co-dependence could help to explain why depletion of distinct nucleolar proteins cause similar downstream cellular phenotypes. Future studies will be needed to mechanistically understand how RNA and protein levels of nucleolar proteins are co-regulated and maintain their functional localization.

### Prolonged nucleolar defects cause profound cytoplasmic re-organization

Interestingly, our unbiased genome-wide CRISPRi screen identified almost 400 non-essential genes which functionally interact with Fibrillarin, providing a comprehensive functional network revealed by nucleolar defects. While synthetic-lethality is observed with components involved in ribosome biosynthesis, many gene deletions rescue survival and/or proliferation of FBL-depleted cells. Surprisingly, these functional suppressors include multiple genes regulating Golgi expansion and vesicular transport, uncovering a previously unknown link between nucleolar stress and cytoplasmic organelle homeostasis. Most prominently scores TP53, which critically regulates both cell cycle progression and cytoplasmic alterations. While the molecular mechanisms of how TP53 executes cytoplasmic reorganization and secretion remain unclear, depletion of the cell cycle inhibitor CDKN1A alone is sufficient to restrict Golgi expansion, possibly by increased Golgi turnover during mitosis. Conversely, cytoplasmic processes are suppressed when abrogating NuGETS, suggesting that an enlarged Golgi and upregulated endo-lysosomal vesicles negatively regulate cell proliferation, possibly by promoting cellular quiescence. For example, RNAi depletion of the adaptor protein TOM1 that is involved in membrane trafficking of ubiquitinated proteins, regulating autophagy, ubiquitination-dependent signaling and endosomal receptor recycling pathways^58,59^ reduces Golgi and endosomal expansion in FBL-depleted cells. We conclude that persistent nucleolar defects trigger a TP53 dependent cytoplasmic adaptation response which promotes cell cycle exit and alters cell-fate into quiescence.

### Prolonged nucleolar defects activate TP53, which in turn alters nucleolar function

Co-depletion of FBL and TP53 depletion rescues cytoplasmic re-organization, but surprisingly also reverts the one-nucleolar morphology. This indicates that TP53 not only senses nucleolar defects but also regulates its function. TP53 activation involves components of the 5S RNP complex including RPL5 and RPL11, but TP53 stabilization and CDKN1A expression levels appear weaker upon FBL-depletion compared to cells exposed to acute stress conditions. Clearly, loss of other ribosomal proteins and 5S RNA also contribute to TP53 activation upon ribotoxic stress^77,79–84^. Interestingly, RPL5 or RPL11 depletion leads to a distinct nucleolar morphology with an enlarged GC phase, occupying 60%-80% of the nucleolar volume. This nucleolar phenotype is associated with DNA damage-mediated TP53 activation and cellular senescence^47^. In contrast, co-depletion of FBL with the 5S RNP components RPL5, RPL11 and RPF2 maintained the one-nucleolar morphology, implying that the nucleolar defects arising from FBL depletion are distinct and occur upstream of the 5S RNP function in nucleolar ribosome biogenesis. Thus, additional pathways must exist that link nucleolar morphology and TP53 levels.

Of note, more than 50% of human cancers are reported to harbor TP53 mutations, which may explain discrepancies observed in nucleolar number and morphology, and different stress responses among cancerous and non-cancerous cell lines^85^. While non-cancerous cell lines like MCF10A robustly change the nucleolar number upon loss of nucleolar proteins, HeLa cells show no or minimal changes^17,77^. Many cancer cells exhibit adaptations to tolerate defective ribosome biogenesis and continue proliferation to sustain increased demands of protein production^47,86,87^. Interestingly, previous studies uncovered a role for TP53 in mediating secretory phenotypes in the context of cancer progression in cell lines derived from cancerous tissues. The overexpression of mutant TP53 or even loss of TP53 function lead to pro-metastatic phenotypes via upregulation of Golgi and secretory pathways^88,89^. While the underlying reason for this discrepancy remains to be explored, it has been hypothesized that prolonged TP53 activation under conditions of low ribosome biogenesis or translation activity originating from loss of function mutations in ribosomal proteins selects for TP53 mutations^29,90^. These mutations may help cells to activate secondary signaling pathways to promote proliferation despite reduced ribosome biogenesis^47^.

### Cells exposed to prolonged nucleolar defects resemble a partial EMT phenotype

Collectively, the expression and functional data of RPE1 cells lacking a functional nucleolus suggest that the cells adapt properties characteristic for an epithelial-to-mesenchymal transition (EMT), manifested by increased secretion and cellular motility. This reversible and plastic embryonic developmental program helps to establish fully differentiated and specialized cells to form functional three-dimensional organs^91,92^. The EMT transition results from changes in transcriptional patterns causing loss of epithelial characteristics and gain in mesenchymal traits. Specialized EMT programs can also be activated by fully differentiated cells leading to wound healing, tissue regeneration and fibrosis as well as metastatic transformation in cancer cells^93,94^. Cancer cells often undergo partial EMT or EM plasticity as they showcase mixed traits of both epithelial and mesenchymal cell types^95,96^. Cellular EMT phenotypes include loss in cell-cell contacts and adhesions, change in cell morphology, accumulation of stress fibers, gain in invasive mobility and upregulated secretion, relying on ECM and secreted ECM components^97^. The cells cease proliferation and increase their survival by adapting a quiescent state, characterized by activating TGFB2 or p38a signaling pathways^93^. The transcription factor SNAI1/2 dominates cell survival signaling ^98–100^. SNAI2 mediated EMT is often driven by c-MYB activation, but MYB and MYBBP1A expression is downregulated in cells with prolonged nucleolar defects^101,102^. Interestingly, TP53 up-regulates some EMT components such as TGFB2, and the 5S RNP complex subunit RPF2 triggers EMT phenotypes via SNAI1/2^103,104^. Thus, TP53 activation via 5S RNP complexes may link nucleolar stress to the observed EMT-like cellular phenotypes. pEMT phenotypes are associated with increased invasive reprogramming and are clinically relevant for patient prognosis^105^. Given this striking resemblance, it is intriguing to speculate that the discovered NuGETS pathway triggered by persistent nucleolar defects may link ribosomopathies and cancer progression. Our results thus open the possibility to identify different classes of cooperative mutations occurring during the hypo-proliferation phase of ribosomopathies, which may facilitate entry into cell cycle. Such cooperative mechanisms could provide attractive therapeutic targets to tackle rare and therapy-resistant malignancies.

**Table 1:** Antibodies.

| Antibodies |  |  |
| --- | --- | --- |
| Mouse monoclonal anti-Nucleophosmin [FC82291] | Abcam | AB10530 |
| Rabbit Recombinant anti-Nucleophosmin [SP236] | Abcam | ab183340 |
| Rabbit Anti-Fibrillarin | Cell Signaling | 2639S |
| Rabbit Anti-Fibrillarin | Abcam | ab5821 |
| Rabbit polyclonal Anti-UTP18 | Invitrogen | PA5-90468 |
| Rabbit Anti-UTP18 | Abcam | ab241578 |
| Mouse Anti-UBF (F-9) | Santa Cruz Biotechnology | sc-13125 |
| Rabbit Anti-BYSL | Abcam | ab251811 |
| Mouse RRP12 (A-3) | Santa Cruz Biotechnology | sc-398593 |
| Rabbit monoclonal RPB1 NTD (D8L4Y) | Cell signaling | 14958 |
| Rabbit Anti-RNA Pol II CTD repeat YSPTSPS (phospho S2) | Abcam | ab5095 |
| Rabbit Anti-RNA Pol II CTD repeat YSPTSPS (phospho S5) [4H8] | Abcam | ab5408 |
| Mouse Anti-RPS5 (A-8) | Santa Cruz Biotechnology | sc-390935 |
| Mouse Anti-RPS3 (C-7) | Santa Cruz Biotechnology | sc-370008 |
| Mouse Anti-p53 (DO-1) | Santa Cruz Biotechnology | sc-126 |
| Mouse Anti-CDKN1A p21 (F-5) | Santa Cruz Biotechnology | sc-6246 |
| Mouse monoclonal Anti-GAPDH | Sigma Fluka Aldrich | G8795 |
| Mouse Purified Anti-GM130 | BD Biosciences | 610823 |
| Mouse IgG1 Anti-EEA1 Clone 14 | BD Transduction Laboratories | AB_397830 |
| Rabbit Anti-TFEB | Abcam | ab181342 |
| Mouse Anti- LAMP1 (D4O1S) | Cell Signaling | 15665S |
| Mouse Anti-GORASP1 (D-12) | Santa Cruz Biotechnology | sc-374423 |
| Rabbit polyclonal Anti-GORASP2 | Protein Tech Group | 10598-1-AP |
| Mouse monoclonal Anti-GORASP2 1C9A3 | Protein Tech Group | 66627-1 |
| Rabbit Recombinant Monoclonal Anti-COL4A1 (HL1351) | Invitrogen | MA5-47009 |
| Rabbit Polyclonal Anti-Fibronectin | Protein Tech Group | 15613-1-AP |
| Mouse Monoclonal Anti-Vimentin | Sigma Fluka Aldrich | V2258 |
| Mouse Monoclonal Anti-Alpha tubulin | Sigma Fluka Aldrich | T5168 |

**Table 2:**
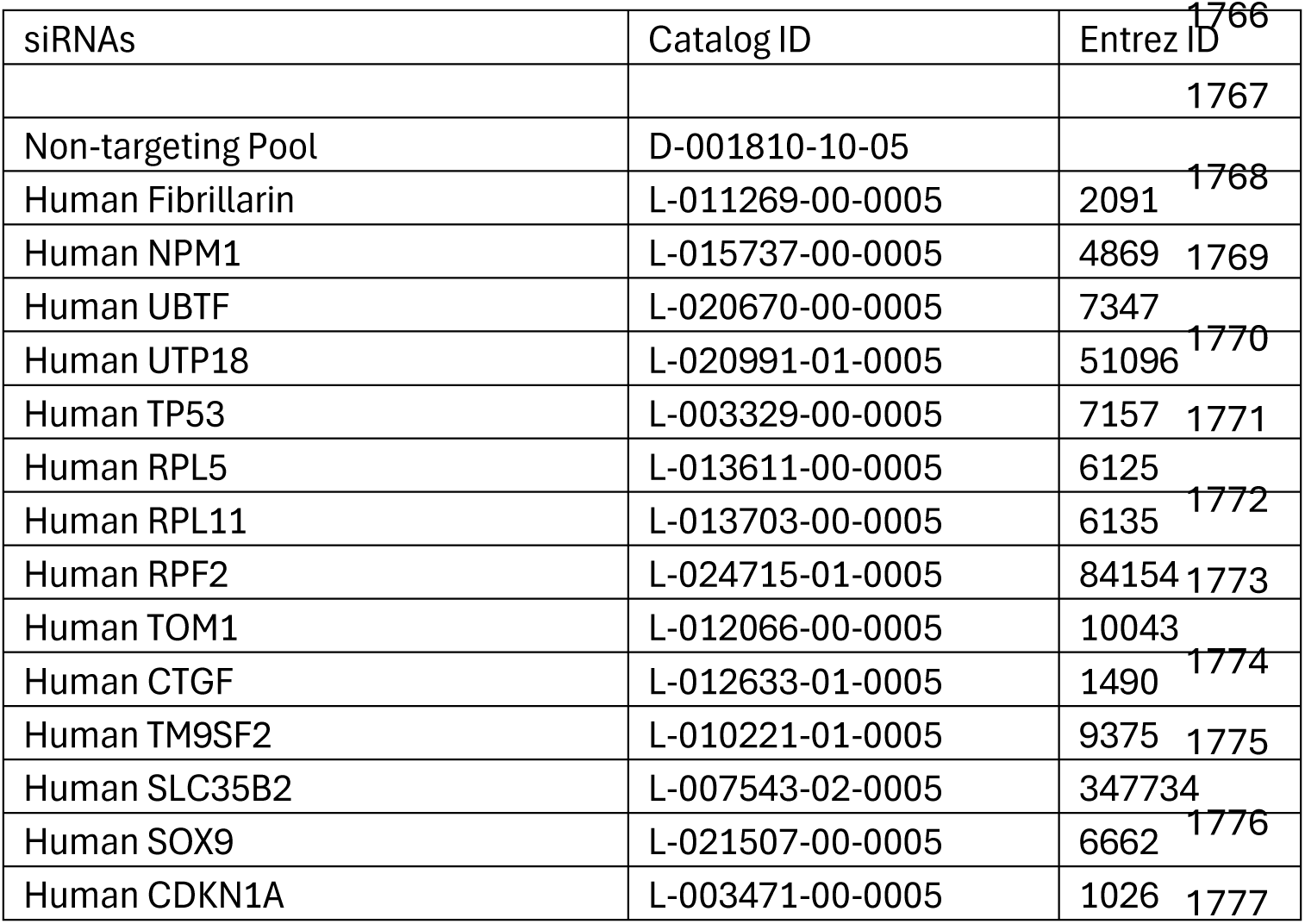
siRNAs.

**Table 3:**
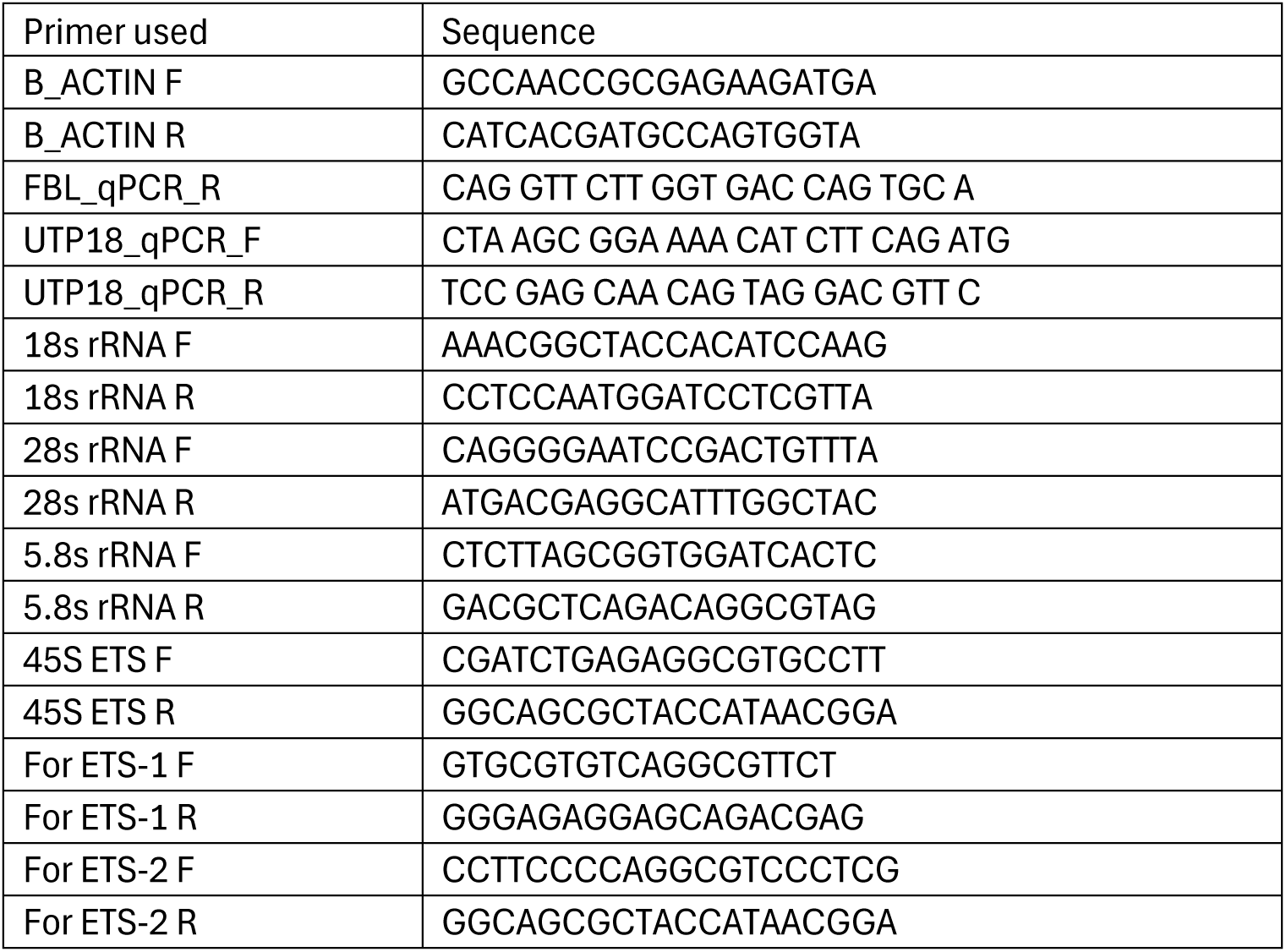
Primers.

## Acknowledgements

We thank Nicole J. Machi from ScopeM for helping with sample preparation and image acquisition of TEM experiments, and Sandhya Manohar for helping with Mass spectrometry sample preparation. We are grateful to John Fielden for kindly providing the RPE1 dCas9-KRAB cell line, Maria Domenica Moccia, Lennart Opitz and the Functional Genomics Center Zürich (FGCZ) platform for sequencing experiments and analysis. We thank Eliana Bianco, Martina Bonassera, Yuriy Rahm and members of Peter laboratory for valuable scientific input, and Alicia Smith for editing the manuscript. This work benefited from support from the ETH technology platforms ScopeM and FGCZ, and the staff of the Institute of Biochemistry. Z.K. acknowledges support from ETH Zurich, which enabled his contributions to this study. J.E.C. is funded by the NOMIS Foundation, the Lotte and Adolf Hotz-Sprenger Stiftung, the Swiss National Science Foundation (SNSF) (project grants 188858, 227979, and 201160), and the European Research Council (ERC) under the European Union’s Horizon 2020 research and innovation program (grant agreement No 855741, DDREAMM). This work was supported by the post-doctoral fellowship EMBO ALTF 627_2021 awarded to P.R., and project grants from the SNSF (project grants 208188 and 200426), and ETH Zürich.

## Author contributions

P.R. and M.P. conceptualized the study. P.R. designed, executed and interpreted the experiments with assistance of T.Q. under the supervision of M.P. Mass Spec. experiments were run and analyzed by F.U. and I.K. S.S.L executed quantitative phase imaging and analyzed images. Z.K. and J.E.C. provided guidance on the CRISPRi screen design and interpretation and contributed the lentiviral library. P.R., T.Q., and I.K. generated the figures. P.R. and M.P. wrote the manuscript with input from all collaborators and authors.

## Declaration of interests

J.E.C. is a co-founder and SAB member of Serac Biosciences and a SAB member of Relation Therapeutics, Hornet Bio, and Kano Therapeutics. The lab of J.E.C. has had funded collaborations with Allogene, Cimeio, and Serac.

---

**Figure S1:**
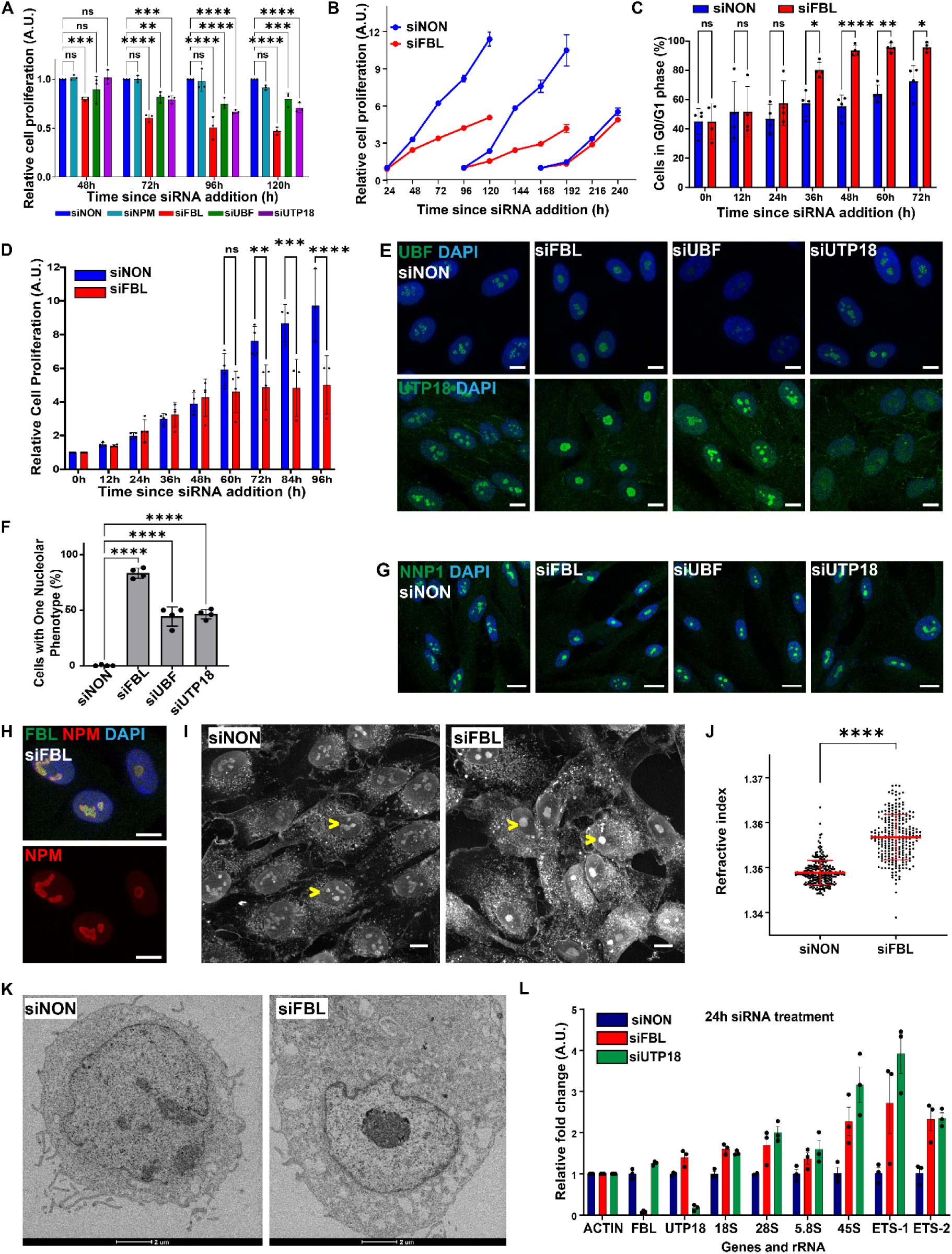
Chronic nucleolar defects preserve the nucleolar proteome but alter their material properties, related to Figure 1. **A.** Bar graph showing cell proliferation of RPE1 cells determined by MTT assay at the indicated time points in hours (h) starting with the addition of the indicated siRNA (siNON, siNPM, siFBL, siUBF, siUTP18). The siNON value was set to 1 for each time-point measurement, and all other values were normalized to siNON. Mean with SD is plotted from N=4 biological replicates. Statistical analysis was performed by a two-way Anova test with Dunnett’s multiple comparison. Asterisks denote p value (****, <0.0001; ***, <0.001; **, <0.01). ns, non-significant. **B.** Line graph showing cell proliferation of RPE1 cells determined by MTT assay over time in hours (h), starting with the addition of siNON and siFBL oligos (time 0). The knockdown cells were replated after 24-hour, 96-hour and 192-hours. The cell number of siNON controls was set to 1 for each time point, and all values from the same plating experiment were normalized respectively. Measurements are from N=4 samples. **C.** Bar graph of data from Fig 1B, quantifying siNON and siFBL treated FUCCI-RPE1 cells in the G0/G1 cell cycle phase at the indicated time points. Mean with SD is plotted from N=4 biological replicates. Statistical analysis was performed by a two-way Anova test with Sidak’s multiple comparison. Asterisks denote p value (****, <0.0001; **, 0.01; *, 0.05). ns, non-significant. **D.** Bar graph of data from Fig 1B, quantifying proliferation of siNON and siFBL treated FUCCI-RPE1 cells determined by MTT assay at the indicated times in hours (h). The 24-hour siRNA value was set to 1, and all other values were normalized accordingly. Mean with SD is plotted from N=3 biological replicates. Statistical analysis was performed by a two-way Anova test with Sidak’s multiple comparison. Asterisks denote p value (****, <0.0001; ***, <0.001; **, <0.01). ns, non-significant. **E.** Representative confocal microscopic images visualize the nucleolus in RPE1 cells treated with the indicated siRNA for 96 hours (siNON, siFBL, siUBF, siUTP18). The top row shows merged co-localization of UBF (green) and DAPI (blue), and the bottom row merged co-localization of UTP18 (green) and DAPI (blue). Images are representative of N=4 biological replicates. Scale bar: 10 µm. **F.** Bar graph showing the percentage (%) of RPE1 cells classified as having 1 nucleolus (indicated as one nucleolar phenotype) at 96 hours siRNA treatment, as observed with NPM staining of cells from Fig 1C. Mean with SD is plotted from N=4 biological replicates. Statistical analysis was performed by a one-way Anova test with Tukey’s multiple comparison. Asterisks denote p value (****, <0.0001). **G.** Representative confocal microscopic images of the nucleolus in RPE1 cells treated with the indicated siRNA for 96 hours (siNON, siFBL, siUBF, siUTP18), and co-stained for NNP-1 (green) and DAPI (blue). Images are representative of N=3 biological replicates. Scale bar: 10 µm. **H.** Representative confocal microscopic images of the nucleolus in RPE1 cells treated with siFBL oligos for 192 hours, and co-stained for NPM (red), FBL (green) and DAPI (blue). Images are representative of N=2 biological replicates. Scale bar: 10 µm. **I.** Representative quantitative phase images (QPI) of RPE1 cells treated with siNON or siFBL oligos for 96 hours. Arrows mark nucleoli. Images are representative of N=3 biological replicates. Scale bar: 10 µm. **J.** Dot plot of RPE1 cells from Fig S1I showing the refractive index (RI) of nucleoli measured by QPI 96 hours after siRNA addition. Mean with SD is plotted from N=3 biological replicates. Statistical analysis was performed by an unpaired t-test. Asterisk denotes the p value (****, <0.0001). **K.** Transmission electron micrographs showing the nuclear morphology of RPE1 cells treated with siNON and siFBL oligos for 96 hours. NE: nuclear envelope. Scale bar: 2 µm. **L.** Bar graph showing the fold change of transcripts corresponding to the indicated genes and ribosomal RNA species detected by RT-qPCR performed on cDNA derived from siNON-, siFBL- and siUTP18-depleted RPE1 cells at 24 hours post siRNA treatment. Actin RNA levels in siNON treated cells were set to 1, and all other values were normalized. Mean with SD is plotted from N=3 biological replicates.

**Figure S2:**
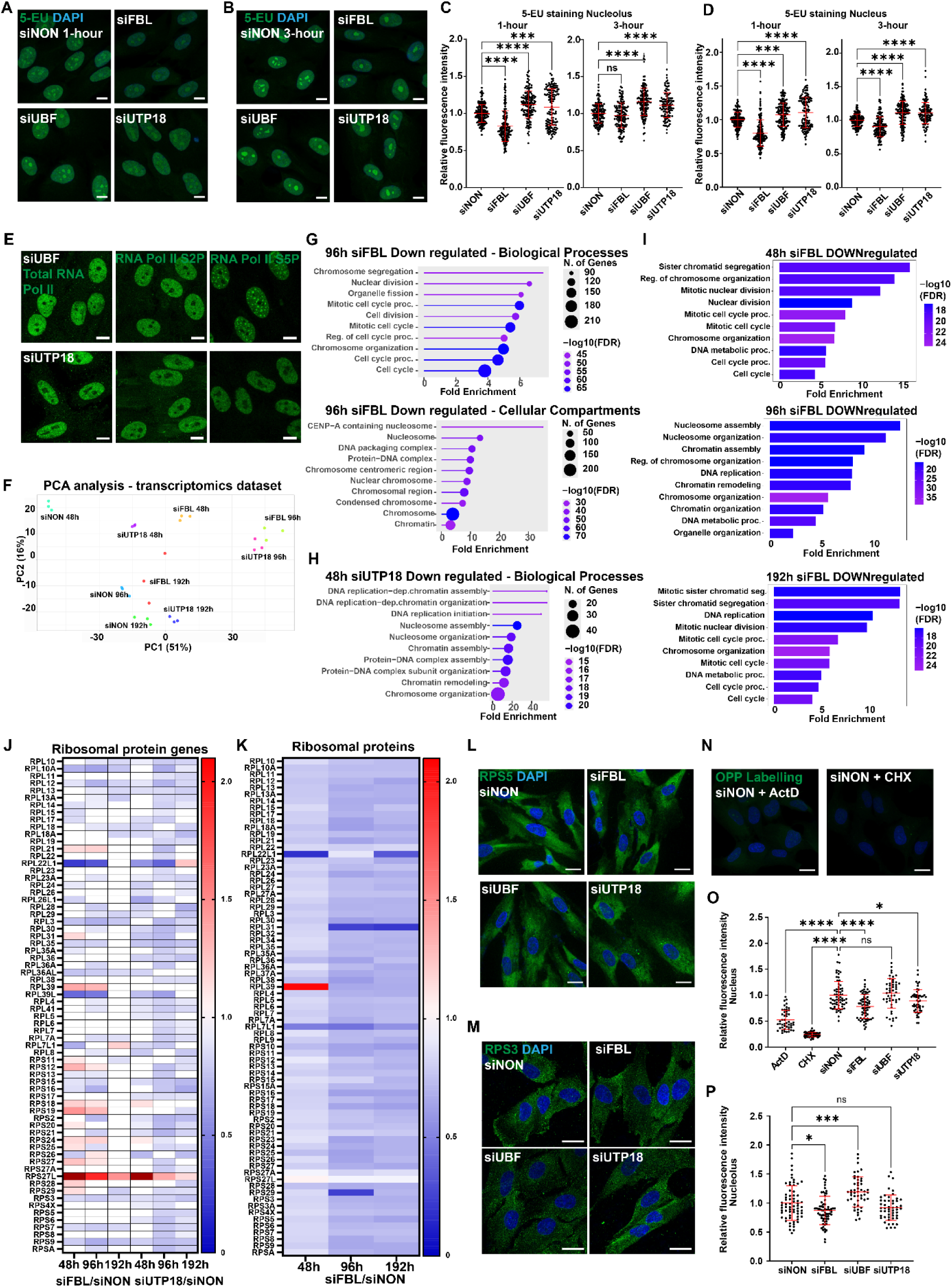
Pol I and Pol II transcription and cytoplasmic translation remain active in cells with loss of nucleolar morphology and ribosome biogenesis defects, Related to Figure 2. **A.** Representative confocal microscopic images show RPE1 cells treated with the indicated siRNA for 96 hours (siNON, siFBL, siUBF, siUTP18), incubated for 1 hour with 1 mM 5-ethynyl-uridine (5-EU) and co-stained with AF488-Azide (green) and DAPI (blue). Images are representative of N=4 biological replicates. Scale bar: 10 µm. **B.** Representative confocal microscopic images showing RPE1 cells treated with the indicated siRNA for 96 hours (siNON, siFBL, siUBF, siUTP18), incubated with 1 mM 5-EU for 3 hours and co-stained for 5-ethynyl-uridine (5-EU) with AF488-Azide (green) and DAPI (blue). Images are representative of N=4 biological replicates. Scale bar: 10 µm. **C.** Dot plot showing data from Fig S2A-B, measuring the fluorescence intensity for 5-EU with AF488-Azide in the nucleolus after 1 or 3 hours 5-EU addition following treatment of RPE1 cells with the indicated siRNA for 96 hours (siNON, siFBL, siUBF, siUTP18). All values were normalized to the siNON average of the corresponding experiment. The mean with SD is plotted from N=3 biological replicates. Statistical analysis was performed by an ordinary one-way Anova test with Dunnett’s multiple comparison. Asterisks denote p value (****, <0.0001). ns, non-significant. **D.** Dot plot showing data from Fig S2A-B, measuring the fluorescence intensity for 5-EU with AF488-Azide in the nucleus for 1 or 3 hours 5-EU addition following treatment of RPE1 cells with the indicated siRNA for 96 hours (siNON, siFBL, siUBF, siUTP18). All values were normalized to siNON average of corresponding experiment. The mean with SD is plotted from N=3 biological replicates. Statistical analysis was performed by an ordinary one-way Anova test with Dunnett’s multiple comparison. Asterisks denote p value (****, <0.0001). **E.** Representative confocal microscopic images of RPE1 cells treated with siUBF (top row) or siUTP18 (bottom row) for 96 hours, and stained for total RNA Polymerase II (green, left), or serine-2 (green, middle) or serine 5 (green, right) phosphorylation of the C-terminal tail of RNA Polymerase II. Images are representative of N=3 biological replicates. Scale bar: 10 µm. **F.** Principal component analysis (PCA) for transcriptomic (RNA-seq) datasets for the top 5000 features ranked by S.D. The data is plotted for normalized log2 counts using 2 principal components with highest variability (PC1 = 51%, PC2 = 16%) using FGCZ Shiny Apps. **G.** Lollipop chart of gene enrichment analysis of RNA-seq datasets from RPE1 cells treated with siFBL or siNON for 96 hours using ShinyGO for genes with more than 1.5-fold downregulation and a p-value cutoff of 0.05. The top panel represents biological processes, and the bottom panel cellular component enrichment. Pathways over fold enrichment were plotted with -log10FDR using the indicated color scheme. Number of genes: 622 and FDR cutoff: 0.01. **H.** Lollipop chart of biological processes enrichment analysis of RNA-seq datasets from RPE1 cells treated with siUTP18 or siNON for 48 hours using ShinyGO for genes with more than 1.5-fold downregulation and a p-value cutoff of 0.05. Pathways over fold enrichment were plotted with -log10FDR using the indicated color scheme. Number of genes: 146 and FDR cutoff: 0.01. **I.** Bar plot chart of biological processes enrichment analysis of mass-spectrometry datasets prepared from RPE1 cells treated with siFBL or siNON controls for 48 (top panel), 96 (middle panel) and 192 hours (bottom panel) using ShinyGO for genes with more than 1-fold downregulation and a p-value cutoff of 0.05. Pathways over fold enrichment were plotted with -log10FDR using the indicated color scheme. Number of genes: 137, 288 and 187 respectively and FDR cutoff: 0.01. **J.** Heat map representing gene fold change of ribosomal proteins from RNA-seq data from RPE1 cells treated with siFBL, siUTP18 or siNON controls for 48-, 96- and 192 hours. The scale bar ranges from 0 to 2-fold change and genes follow a p-value cutoff of 0.05. Out of scale values are represented in maroon. Non-significant values are left blank and represented in white. The gene list is based on RNAseq data using the RPL and RPS nomenclature. **K.** Heat map representing protein level fold change of ribosomal proteins from RNA-seq data of RPE1 cells treated with siFBL, siUTP18 or siNON controls for 48, 96 and 192 hours. The scale bar ranges from 0 to 2-fold change and proteins follow a p-value cutoff of 0.05. Out of scale values are represented in maroon. Non-significant values are left blank and represented in white. The protein list is based on proteins identified in MS using the RPL and RPS nomenclature. **L.** Representative confocal microscopic images of RPE1 cells treated with the indicated siRNA for 96 hours (siNON, siFBL, siUBF, siUTP18), and co-stained with RPS5 (green) and DAPI (blue). Images are representative of N=3 biological replicates. Scale bar: 10 µm. **M.** Representative confocal microscopic images of RPE1 cells treated with the indicated siRNA for 96 hours (siNON, siFBL, siUBF, siUTP18), and co-stained for RPS3 (green) and DAPI (blue). Images are representative of N=2 biological replicates. Scale bar: 20 µm. **N.** Representative confocal microscopic images of RPE1 cells treated with 625nM Actinomycin D (ActD) or 30µg/ml Cycloheximide (CHX) for 2 hours, and co-stained for O-Propargyl-puromycin with AF488-Azide (green) and DAPI (blue). 10 µM OPP was added for 3 hours. Images are representative of N=3 biological replicates. Scale bar: 10 µm. **O.** Dot plot showing data from Fig 2K and S2N, measuring the fluorescence intensity for O-Propargyl-puromycin with AF488-Azide in the nucleus of RPE1 cells treated with the indicated siRNA for 96 hours (siNON, siFBL, siUBF, siUTP18) and 625nM Actinomycin D (ActD) and 30µg/ml Cycloheximide (CHX) for 2 hours. All values were normalized to siNON average of corresponding experiments. The mean with SD is plotted from N=3 (siRNA) and N=2 (inhibitors) biological replicates. Statistical analysis was performed by an ordinary one-way Anova test with Dunnett’s multiple comparison. Asterisks denote p value (****, <0.0001; *, <0.05). ns, non-significant. **P.** Dot plot showing data from Fig 2K and S2N, measuring the fluorescence intensity for O-Propargyl-puromycin with AF488-Azide in the nucleolus of RPE1 cells treated with the indicated siRNA for 96 hours (siNON, siFBL, siUBF, siUTP18). All values were normalized to siNON average of corresponding experiments. The mean with SD is plotted from N=3 biological replicates. Statistical analysis was performed by an ordinary one-way Anova test with Dunnett’s multiple comparison. Asterisks denote p value (***, <0.001; *, <0.05). ns, non-significant.

**Figure S3:**
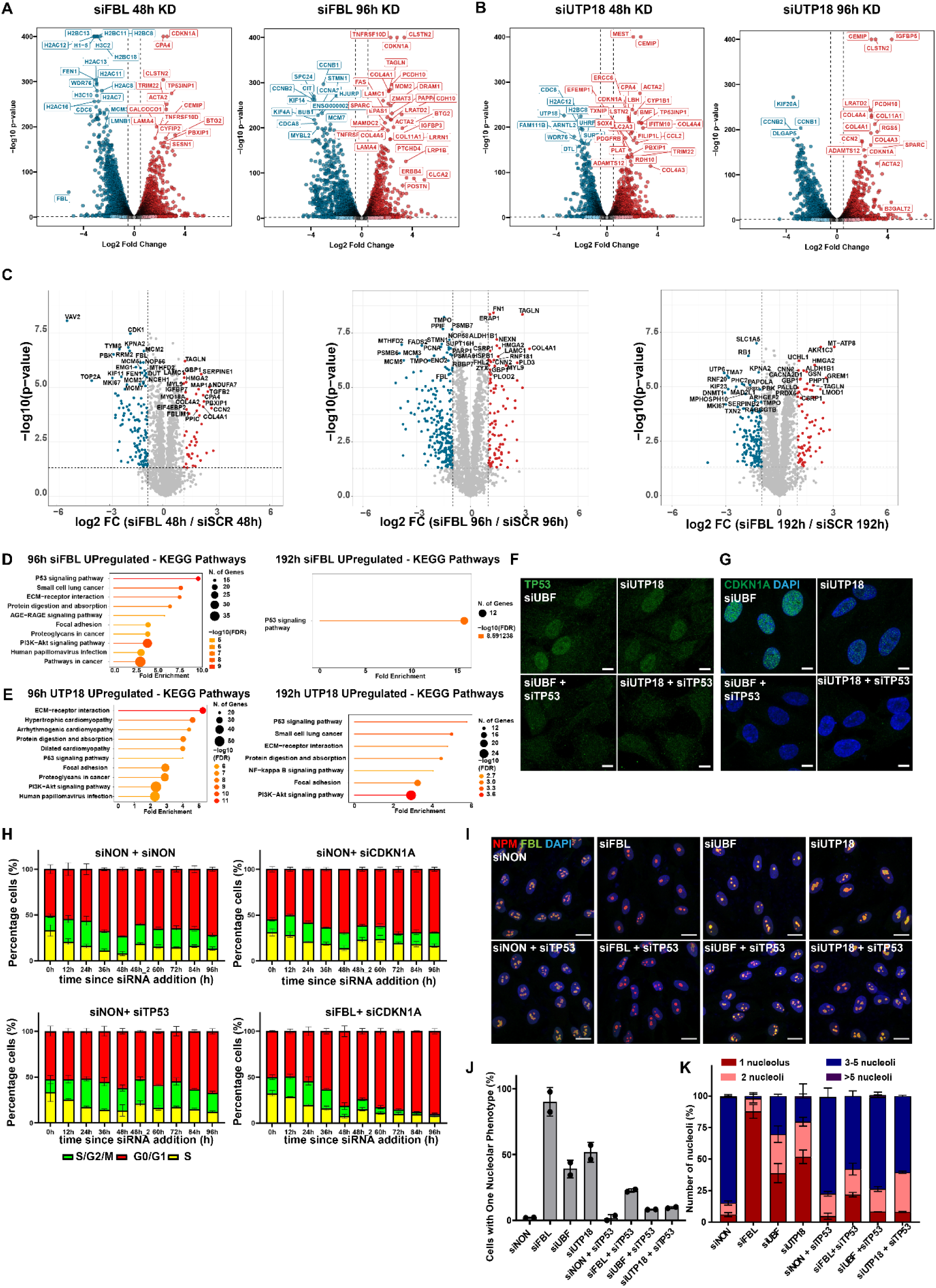
Co-depletion of TP53 rescues cell cycle and nucleolar morphology defects associated with loss of nucleolar proteins, related to Figure 3. **A.** Volcano Plot showing differentially expressed genes from the RNA-seq analysis in RPE1 cells treated with siFBL and siNON controls for 48 (left panel) or 96 hours (right panel). Significant genes in siFBL over siNON are marked with log_2_ fold change > 1 and an adjusted p value (p adj) < 0.05. Downregulated genes upon FBL depletion are represented in blue and upregulated genes in red. The volcano plots are generated using FGCZ Shiny Apps. **B.** Volcano Plot showing differentially expressed genes from the RNA-seq analysis in RPE1 cells treated with siUTP18 and siNON controls for 48 (left panel) or 96 hours (right panel). Significant genes in siUTP18 over siNON are marked with log_2_ fold change > 1 and an adjusted p value (p adj) < 0.05. Downregulated genes upon FBL depletion are represented in blue and upregulated genes in red. The volcano plots are generated using FGCZ Shiny Apps. **C.** Volcano Plot showing differentially expressed proteins from the total proteome mass-spectrometry analysis of RPE1 cells treated with siFBL and siNON controls for 48 (left panel), 96 (middle panel) or 192 hours (right panel). Proteins that significantly differ in siFBL and siNON controls are marked with log_2_ fold change > 1 and an adjusted p value (p adj) < 0.05. Downregulated proteins in FBL depleted cells are represented in blue and upregulated proteins in red. The volcano plots are generated using R and top differentially expressed proteins are labelled with no specific criteria. **D.** Lollipop chart of KEGG (Kyoto encyclopedia of genes and genomes) enrichment analysis of RNA-seq datasets from Fig S3A generated from RPE1 cells treated with siFBL or siNON controls for 96 (left panel) or 192 hours (right panel) using ShinyGO to mark genes with more than 2-fold upregulation and a p-value cutoff of 0.05. Pathways over fold enrichment were plotted with -log10FDR using the indicated color scheme. Number of genes: 547 and 237 respectively and FDR cutoff: 0.01. **E.** Lollipop chart of KEGG (Kyoto encyclopedia of genes and genomes) enrichment analysis of RNA-seq datasets from Fig S3B prepared from RPE1 cells treated with in siUTP18 or siNON controls for 96 (left panel) or 192 hours (right panel) using ShinyGO to identify genes with more than 2-fold upregulation and a p-value cutoff of 0.05. Pathways over fold enrichment were plotted with -log10FDR using the indicated color scheme. Number of genes: 691 and 597 respectively and FDR cutoff: 0.01. **F.** Representative confocal microscopic images acquired as in Fig 3D showing RPE1 cells treated with the indicated single or double siRNAs for 96 hours (siNON, siUBF, siUTP18, siUBF+siTP53, siUTP18+siTP53), and co-stained for TP53 (green) and DAPI (blue). Images are representative of N=4 biological replicates. Scale bar: 10 µm. **G.** Representative confocal microscopic images acquired as in Fig 3E showing RPE1 cells treated with the indicated single or double siRNAs for 96 hours (siNON, siUBF, siUTP18, siUBF+siTP53, siUTP18+siTP53), and co-stained for CDKN1A (green) and DAPI (blue). Images are representative of N=4 biological replicates. Scale bar: 10 µm. **H.** Bar graph showing the percentage (%) of FUCCI-RPE1 cells in different cell cycle phases (S = yellow; S/G2/M = green; G0/G1=red) at the indicated time in hours (h) relative to the siNON + NON (top left panel), siNON + siTP53 (bottom left panel), siNON + siCDKN1A (top right panel) or siFBL + siCDKN1A (bottom right panel) double siRNA treatments. The cells were freshly re-plated to avoid confluency at 48h indicated as 48_2. The mean with SD is plotted from N=3 biological replicates. **I.** Representative confocal microscopic images depicting RPE1 cells treated with the indicated single (siNON, siFBL, siUBF, siUTP18) or double (siNON + siTP53, siFBL + siTP53, siUBF + siTP53, siUTP18 + siTP53) siRNA for 96 hours, co-stained for NPM (red), FBL (green) and DAPI (blue). Images are representative of N=4 biological replicates. Scale bar: 20 µm. **J.** Bar graph showing data from Fig S3I, measuring the percentage (%) of RPE1cells classified as having 1 nucleolus (indicated as one nucleolar phenotype) based on NPM staining, following treatment with the indicated single (siNON, siFBL, siUBF, siUTP18) or double (siNON + siTP53, siFBL + siTP53, siUBF + siTP53, siUTP18 + siTP53) siRNA for 96 hours. The mean with SD is plotted from N=2 biological replicates. **K.** Bar graph showing data from Fig S3I, measuring the percentage (%) of RPE1 cells classified as having 1 (maroon), 2 (pink), 3-5 (navy blue) or >5 (purple) nucleoli following treatment with the indicated single (siNON, siFBL, siUBF, siUTP18) or double (siNON + siTP53, siFBL + siTP53, siUBF + siTP53, siUTP18 + siTP53) siRNA for 96 hours, as observed with NPM staining. The mean with SEM is plotted from N=2 biological replicates.

**Figure S4:**
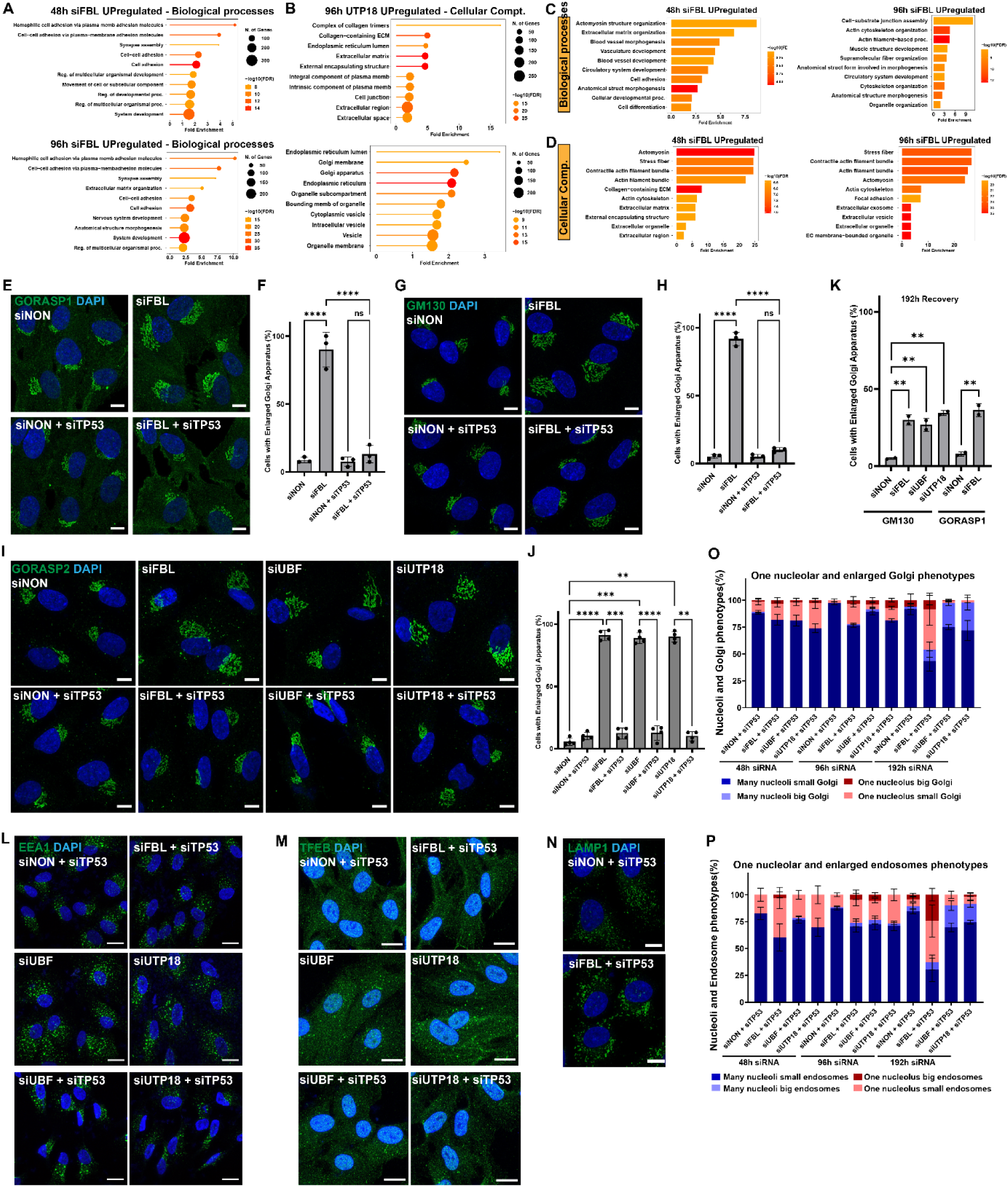
TP53 signaling induced upon chronic nuclear defects rewires organellar homeostasis, related to Figure 4. **A.** Lollipop chart of biological processes enrichment analysis of RNA-seq datasets obtained from RPE1 cells treated with siFBL or siNON controls for 48 (top panel) and 96 hours (bottom panel) using ShinyGO selecting genes with more than 2-fold upregulation and a p-value cutoff of 0.05. Pathways over fold enrichment were plotted with -log10FDR using the indicated color scheme. Number of genes: 978 and 547 respectively and FDR cutoff: 0.01. **B.** Lollipop chart of cellular component enrichment analysis from RNA-seq datasets from RPE1 cells treated with siUTP18 and siNON controls for 96 hours using ShinyGO selecting for genes with more than 2-fold (top panel) and 1.5-fold (bottom panel) upregulation and a p-value cutoff of 0.05. Pathways over fold enrichment were plotted with log10FDR using the indicated color scheme. Number of genes: 691 and 797 respectively and FDR cutoff: 0.01. **C.** Bar plot chart of biological processes enrichment analysis of mass-spectrometry datasets of RPE1 cells treated with siFBL or siNON controls for 48 (left panel) and 96 hours (right panel), analyzed using ShinyGO to select proteins with more than 1.5-fold upregulation and a p-value cutoff of 0.05. Pathways over fold enrichment were plotted with log10FDR using the indicated color scheme. Number of genes: 99 and 256 respectively and FDR cutoff: 0.01. **D.** Bar plot chart of cellular compartment enrichment analysis of mass-spectrometry datasets of RPE1 cells treated with siFBL or siNON controls for 48 (left panel) and 96 hours (right panel) using ShinyGO to select proteins with more than 1.5-fold upregulation and a p-value cutoff of 0.05. Pathways over fold enrichment were plotted with -log10FDR using the indicated color scheme. Number of genes: 99 and 256 respectively and FDR cutoff: 0.01. **E.** Representative confocal microscopic images of RPE1 cells treated with single or double siRNAs for 96 hours (siNON, siFBL, siNON+siTP53, siFBL+siTP53), and co-stained for the Golgi stacking protein GORASP1 (green) and DAPI (blue). Images are representative of N=3 biological replicates. Scale bar: 10 µm. **F.** Bar graph showing the percentage (%) of cells classified with enlarged Golgi using manual annotation of RPE1 cells treated with the indicated siRNA for 96 hours (siNON, siFBL, siNON+siTP53, siFBL+siTP53), as observed with GORASP1 staining of cells shown in Fig S4E. Note that co-depletion of TP53 with FBL revealed significant rescue of the enlarged Golgi phenotype. The mean with SD is plotted from N=3 biological replicates. Asterisks denote p value of (****, <0.0001) as calculated by an ordinary one-way Anova test with Tukey’s multiple comparison. ns: not significant. **G.** Representative confocal microscopic images of RPE1 cells treated with the indicated single or double siRNAs for 96 hours (siNON, siFBL, siNON+siTP53, siFBL+siTP53), and co-stained with the cis-Golgi marker GM130 (green) and DAPI (blue). Images are representative of N=3 biological replicates. Scale bar: 10 µm. **H.** Bar graph showing the percentage (%) of cells classified with an enlarged Golgi using manual annotation of RPE1 cells treated with the indicated siRNA for 96 hours (siNON, siFBL, siNON+siTP53, siFBL+siTP53), as observed with GM130 staining of cells shown in Fig S4G. Note that co-depletion of TP53 and FBL results in significant rescue of the enlarged Golgi phenotype. The mean with SD is plotted from N=3 biological replicates. Asterisks denote p value of (****, <0.0001) as calculated by an ordinary one-way Anova test with Tukey’s multiple comparison. ns: non-significant. **I.** Representative confocal microscopic images of RPE1 cells treated with the indicated single (siNON, siFBL, siUBF, siUTP18) or double (siNON + siTP53, siFBL + siTP53, siUBF + siTP53, siUTP18 + siTP53) siRNA for 96 hours, co-stained for the Golgi stacking protein GORASP2 (green) and DAPI (blue). Images are representative of N=4 biological replicates. Scale bar: 10 µm. **J.** Bar graph showing data from Fig S4I measuring the percentage (%) of cells classified with an enlarged Golgi using manual annotation of RPE1 cells treated with the indicated siRNA for 96 hours (siNON, siFBL, siUBF, siUTP18, siNON + siTP53, siFBL + siTP53, siUBF + siTP53, siUTP18 + siTP53), as observed with GORASP2 staining. Note that co-depletion of TP53 with FBL, UBF and UTP18 show significant rescue of the enlarged Golgi phenotype. The mean with SD is plotted from N=4 biological replicates. Statistical analysis was performed by a two-way Anova test with Tukey’s multiple comparison. Asterisks denote p value (****, <0.0001). ns: non-significant. **K.** Bar graph showing the percentage (%) of cells classified with enlarged Golgi using manual annotation of RPE1 cells treated with the indicated siRNA for 192 hours (siNON, siFBL, siUBF, siUTP18), as observed with GM130 and GORASP1 staining of cells shown in Fig 4E. The mean with SD is plotted from N=2 biological replicates. Statistical analysis was performed by an ordinary one-way Anova test with Tukey’s multiple comparison. Asterisks denote p value (**, <0.01). **L.** Representative confocal microscopic images of RPE1 cells treated with the indicated single (siUBF, siUTP18) or double (siNON + siTP53, siFBL + siTP53, siUBF + siTP53, siUTP18 + siTP53) siRNA for 96 hours, co-stained for the endosomal protein EEA1 (green) and DAPI (blue). Corresponding siFBL and siNON controls are shown in Fig 4I. Images are representative of N=2 biological replicates. Scale bar: 20 µm. **M.** Representative confocal microscopic images of RPE1 cells treated with the indicated single (siUBF, siUTP18) or double (siNON + siTP53, siFBL + siTP53, siUBF + siTP53, siUTP18 + siTP53) siRNA for 96 hours, co-stained for the lysosomal protein TFEB (green) and DAPI (blue). Corresponding siFBL and siNON controls are shown in Fig 4J. Images are representative of N=2 or N=3 biological replicates. Scale bar: 20 µm. **N.** Representative confocal microscopic images of RPE1 cells treated with (siNON + siTP53, siFBL + siTP53) siRNA for 96 hours, co-stained for the endosomal-lysosomal protein LAMP1 (green) and DAPI (blue). Corresponding siFBL and siNON controls are shown in Fig 4J. Images are representative of N=3 biological replicates. Scale bar: 10 µm. **O.** Bar graph showing the percentage (%) of cells classified with one nucleolus phenotype (counted as 1 or 2 nucleoli) with a small (comparable to siNON condition) Golgi, many nucleoli (>2 nucleoli) with a small Golgi, one nucleolus with a big (enlarged) Golgi and many nucleoli with a big Golgi using manual annotation of RPE1 cells treated with the indicated siRNA for 48-, 96- and 192 hours (siNON + siTP53, siFBL + siTP53, siUBF + siTP53, siUTP18 + siTP53), as observed by NPM and GM130 staining. The mean with SEM is plotted from N=3 biological replicates. **P.** Bar graph showing the percentage (%) of cells classified with one nucleolus phenotype (counted as 1 or 2 nucleoli) with small (comparable to siNON) endosomes, many nucleoli (>2 nucleoli) with small endosomes, one nucleolus with big (enlarged and many) endosomes and many nucleoli with big endosomes using manual annotation of RPE1 cells treated with the indicated siRNA for 48-, 96- and 192 hours (siNON + siTP53, siFBL + siTP53, siUBF + siTP53, siUTP18 + siTP53), as observed with NPM and EEA1 staining. The mean with SEM is plotted from N=3 biological replicates.

**Figure S5:**
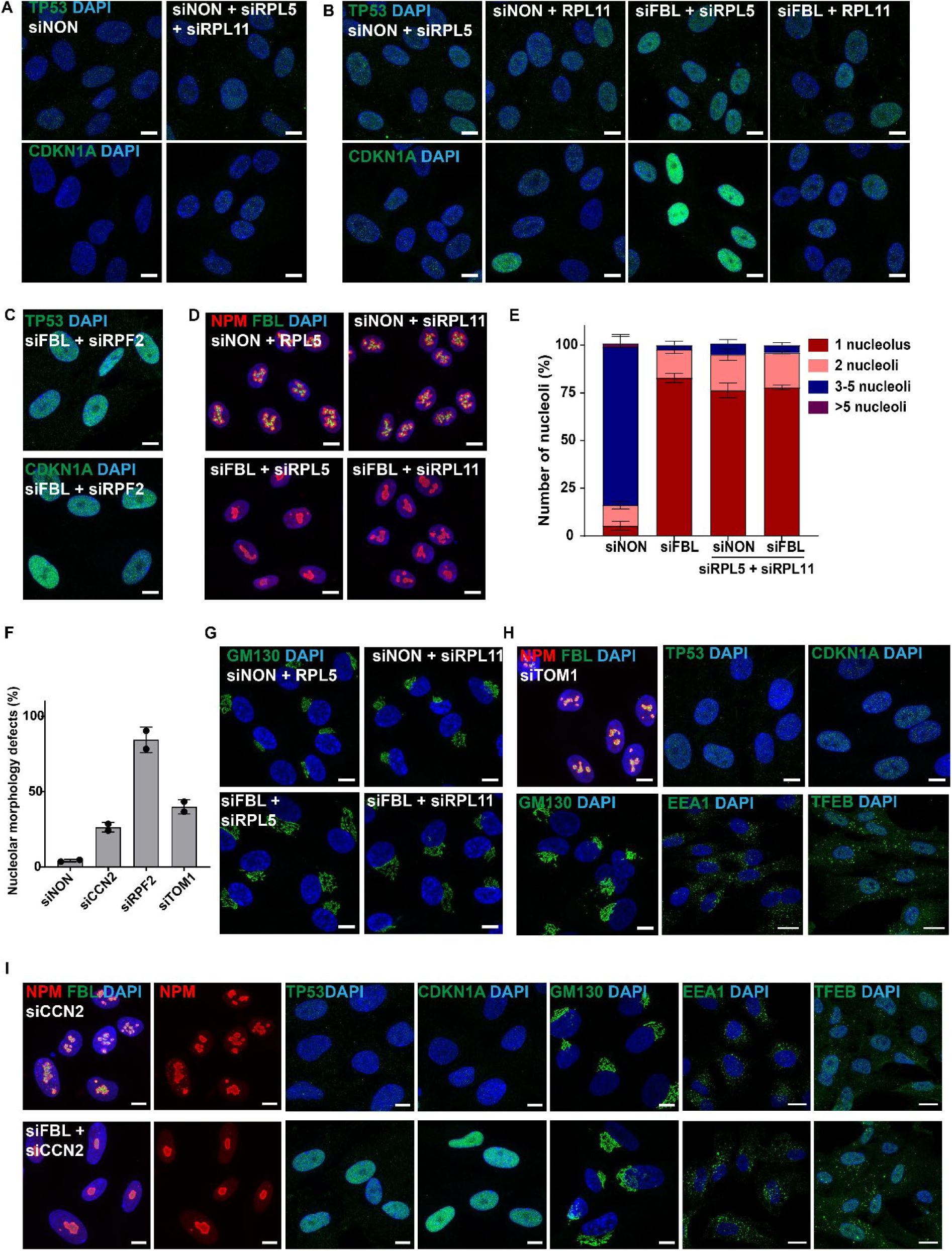
5S RNP dependent activation of TP53 and cytoplasmic rearrangements upon chronic nucleolar defect, related to Figure 5. **A.** Representative confocal microscopic images showing RPE1 cells treated with the indicated single or double siRNAs for 96 hours (siNON, siNON+ siRPL5 + siRPL11). Cells in the top panel are co-stained for TP53 (green) and DAPI (blue), and in the bottom panel for CDKN1A (green) and DAPI (blue). Images are representative of N=2 biological replicates. Scale bar: 10 µm. **B.** Representative confocal microscopic images showing RPE1 cells treated with the indicated single or double siRNAs for 96 hours (siNON + siRPL5, siFBL + siRPL5, siNON + siRPL11, siFBL + siRPL11). Cells in the top panel are co-stained for TP53 (green) and DAPI (blue), and in the bottom panel for CDKN1A (green) and DAPI (blue). Images are representative of N=2 biological replicates. Scale bar: 10 µm. **C.** Representative confocal microscopic images showing RPE1 cells treated with indicated single or double siRNAs for 96 hours (siRPF2). Cells in the top panel are co-stained for TP53 (green) and DAPI (blue), and in the bottom panel for CDKN1A (green) and DAPI (blue). Images are representative of N=2 biological replicates. Scale bar: 10 µm. **D.** Representative confocal microscopic images showing RPE1 cells treated with the indicated siRNA for 96 hours (siNON + siRPL5, siFBL, siRPL5, siNON + siRPL11, siFBL+ siRPL11) and co-stained for NPM (red), FBL (green) and DAPI (blue). Images are representative of N=3 biological replicates. Scale bar: 10 µm. **E.** Bar graph showing data from Fig 5D measuring the percentage (%) of RPE1 cells classified as having 1 (maroon), 2 (pink), 3-5 (navy blue) or >5 (purple) nucleoli, as observed with NPM staining. Note that the nucleolar number classification is independent of nucleolar morphology (irregular/spherical). The mean with SEM is plotted from N=3 biological replicates. **F.** Bar graph showing data from Fig 5 and S5 measuring the percentage (%) of RPE1 cells classified as having aberrant nucleolar morphology and reduced numbers (indicated as nucleolar morphology defects) at 96 hours siRNA treatment, as observed with NPM staining. The mean with SD is plotted from N=2 biological replicates. **G.** Representative confocal microscopic images showing RPE1 cells treated with the indicated siRNA for 96 hours (siNON + siRPL5, siFBL, siRPL5, siNON + siRPL11, siFBL+ siRPL11) and co-stained for the cis-Golgi marker GM130 (green) and DAPI (blue). Images are representative of N=3 biological replicates. Scale bar: 10 µm. **H.** Representative confocal microscopic images showing RPE1 cells treated with TOM1 siRNA for 96 hours and co-stained for NPM (red), FBL (green) and DAPI (blue), TP53 (green) and DAPI (blue) and CDKN1A (green) and DAPI (blue) (top panel, left to right). Cells in the bottom panel (left to right) are co-stained for the cis-Golgi marker GM130 (green) and DAPI (blue), the endosomal marker EEA1 (green) and DAPI (blue) and the lysosomal marker TFEB (green) and DAPI (blue). Images are representative of N=2 biological replicates. Scale bars: 10 µm, 20 µm. **I.** Representative confocal microscopic images showing RPE1 cells treated with siCCN2 (top panel) or siFBL + siCCN2 (bottom panel) for 96 hours and co-stained for (left to right) NPM (red), FBL (green) and DAPI (blue), NPM (red), TP53 (green) and DAPI (blue), CDKN1A (green) and DAPI (blue), the cis-Golgi marker GM130 (green) and DAPI (blue), the endosomal marker EEA1 (green) and DAPI (blue) and the lysosomal marker TFEB (green) and DAPI (blue). Images are representative of N=2 biological replicates. Scale bars: 10 µm, 20 µm.

**Figure S6:**
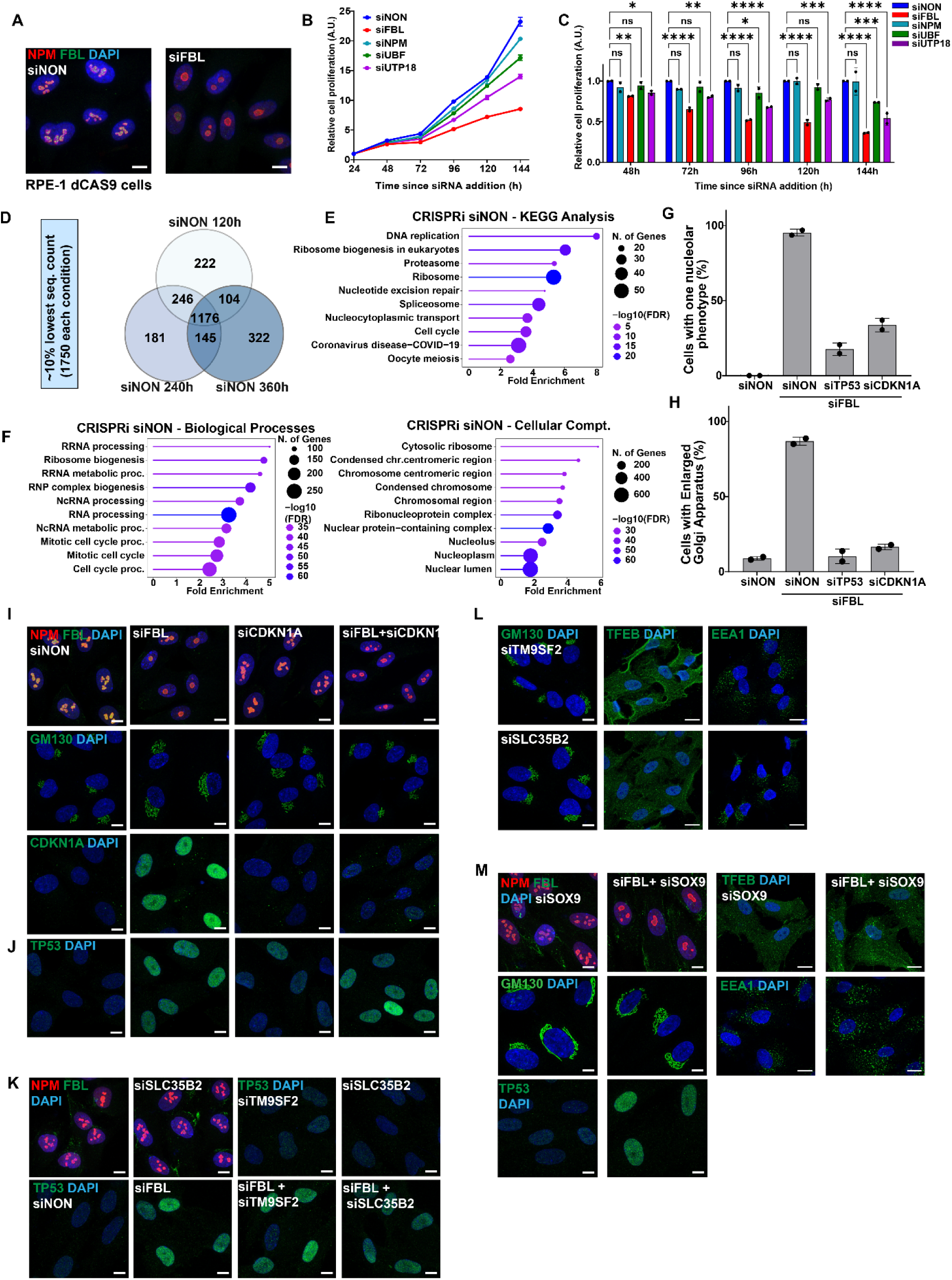
Nucleolar defects associated proliferation rescue by genes identified in CRISPRi screen routes via cellular secretory pathways, related to Figure 6. **A.** Representative confocal microscopic images of RPE1 dCas9-KRAB cells treated with si FBL and siNON control for 96 hours, and co-stained for NPM (red), FBL (green) and DAPI (blue). Images are representative of N=3 biological replicates. Scale bar: 10 µm. **B.** Cell proliferation was determined by MTT assays of RPE1 cells treated with the indicated siRNA (siNON, siNPM, siUBF, siUTP18 and siFBL) at the indicated time points (in hours). The 24-hour siRNA value was set to 1, and all other values were normalized to 24 hours. Measurements are of N=4 samples. **C.** Bar graph showing the cell proliferation determined by MTT assays of RPE1 cells treated with the indicated siRNA for 96 hours (siNON, siNPM, siFBL, siUBF, siUTP18) over time (in hours (h)). The siNON measurement was set to 1 for each time-point, and all other values were normalized to the siNON control. The mean with SD is plotted from N=2 biological replicates. Statistical analysis was performed by a two-way Anova test with Dunnett’s multiple comparison. Asterisks denote p value (****, <0.0001; ***, <0.001; **, <0.01; *, <0.05). ns, non-significant. **D.** Venn Diagram representing overlap of ∼10% genes with the lowest sequencing count in each time point from control siRNA treatment (siNON) identified by deep sequencing of the genome-wide CRISPRi screen. 1750 genes from each condition are picked resulting in a total of 2396 unique genes. 1176 genes overlap in all 3 conditions and 1671 genes in at least two conditions. **E.** Lollipop chart from data in Fig S6D showing KEGG (Kyoto encyclopedia of genes and genomes) enrichment analysis using ShinyGO from the CRISPRi dataset for the 1176 genes common in all 3 conditions. Pathways over fold enrichment were plotted with -log10FDR using the indicated color scheme and circles represent the number of genes. FDR cutoff: 0.01. **F.** Lollipop chart from data in Fig S6D showing biological processes (left panel) and cellular component (right panel) enrichment analysis using ShinyGO from the CRISPRi dataset for the 1176 genes common in all 3 conditions. Pathways over fold enrichment were plotted with -log10FDR using the indicated color scheme and circles represent the number of genes. FDR cutoff: 0.01. **G.** Bar graph showing data from Fig S6I measuring the percentage (%) of cells classified with one nucleolar phenotype of RPE1 cells treated with the indicated siRNA for 96 hours (siNON, siFBL, siFBL + TP53, siFBL + siCDKN1A), as observed with NPM staining. Note that co-depletion of TP53 or CDKN1A and FBL rescue the one nucleolar phenotype. The mean with SD is plotted from N=2 biological replicates. **H.** Bar graph showing data from Fig S6I measuring the percentage (%) of cells classified with an enlarged Golgi phenotype of RPE1 cells treated with the indicated siRNA for 96 hours (siNON, siFBL, siFBL + TP53, siFBL + siCDKN1A), as observed with GM130 staining. Note that co-depletion of TP53 or CDKN1A and FBL rescue the enlarged Golgi phenotype. The mean with SD is plotted from N=2 biological replicates. **I.** Representative confocal microscopic images of RPE1 cells treated with the indicated siRNAs for 96 hours (siNON, siFBL, siFBL + TP53, siFBL + siCDKN1A). The top row displays cells co-stained for the nucleolar proteins NPM (red), FBL (green) and DAPI (blue), the middle row the cis-Golgi protein GM130 (green) and DAPI (blue), and the bottom row for CDKN1A (green) and DAPI (blue). Images are representative of N=2 biological replicates. Scale bar: 10 µm. **J.** Representative confocal microscopic images of RPE1 cells treated with the indicated siRNAs for 96 hours (siNON, siFBL, siFBL + TP53, siFBL + siCDKN1A), and co-stained for TP53 (green) and DAPI (blue). Images are representative of N=2 biological replicates. Scale bar: 10 µm. **K.** Representative confocal microscopic images of RPE1 cells treated with the indicated siRNAs for 96 hours (siNON, siFBL, siTM9SF2, siSLC35B2, siFBL + siTM9SF2, siFBL + siSLC35B2). Cells in the top row (two on left) are co-stained for the nucleolar proteins NPM (red), FBL (green) and DAPI (blue), and the bottom row TP53 (green) and DAPI (blue). The corresponding nucleolar images are part of Fig 6F. Images are representative of N=2 biological replicates. Scale bar: 10 µm. **L.** Representative confocal microscopic images of RPE1 cells treated with the indicated siRNAs for 96 hours (siTM9SF2, siSLC35B2). The left panels show cells co-stained for the cis-Golgi protein GM130 (green) and DAPI (blue), the middle panel co-staining of the lysosomal protein TFEB (green) and DAPI (blue), and the bottom panel of the endosomal protein EEA1 (green) and DAPI (blue). Corresponding images are part of Fig 6I. Images are representative of N=2 biological replicates. Scale bars: 10 µm, 20 µm, 20 µm. **M.** Representative confocal microscopic images of RPE1 cells treated with the indicated siRNAs for 96 hours (siSOX9, siFBL + siSOX9), and co-stained as indicated for the nucleolar proteins NPM (red) and FBL (green), or the lysosomal protein TFEB (green), the cis-Golgi protein GM130 (green), the endosomal protein EEA1 (green), TP53 (green) and DAPI (blue). Images are representative of N=2 biological replicates. Scale bars: 10 µm and 20 µm.

**Figure S7:**
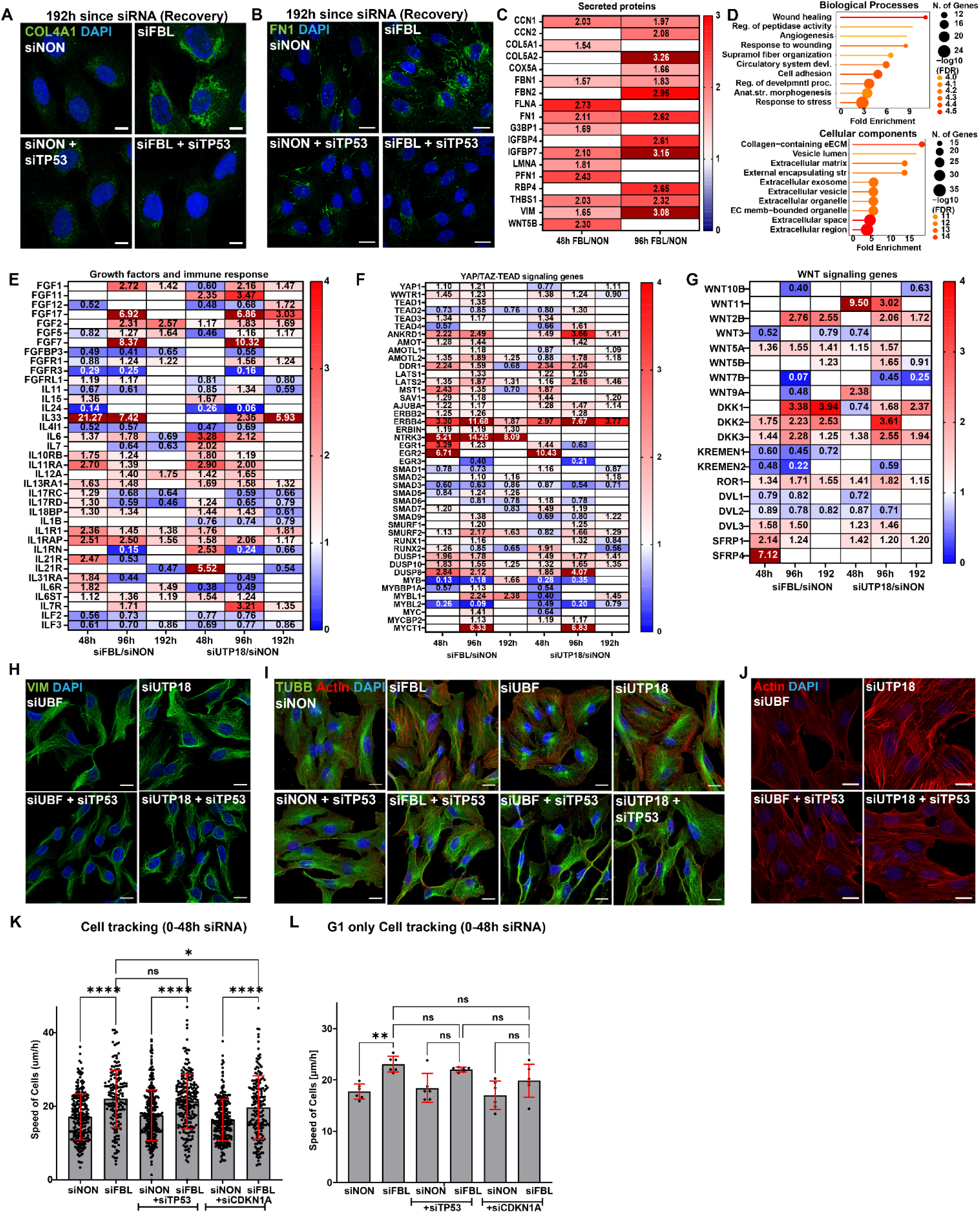
Chronic nucleolar defects trigger cellular quiescence and an EMT like phenotype, related to Figure 7. **A.** Representative confocal microscopic images of RPE1 cells treated with the indicated single or double siRNAs for 192 hours (siNON, siFBL, siNON+siTP53, siFBL+siTP53), and co-stained for collagen COL4A1 (green) and DAPI (blue). Images are representative of N=2 biological replicates. Scale bar: 10 µm. **B.** Representative confocal microscopic images of RPE1 cells treated with the indicated single or double siRNAs for 192 hours (siNON, siFBL, siNON+siTP53, siFBL+siTP53), and co-stained for fibronectin FN1 (green) and DAPI (blue). Images are representative of N=2 biological replicates. Scale bar: 10 µm. **C.** Heat map representing protein level fold change of prominent secreted proteins from cell culture supernatants (secretome) of RPE1 cells treated with siFBL and siNON control for 48- and 96 hours identified by mass spectrometry. The cells were cultured with fetal calf serum (FCS) prior to collection. The scale bar ranges from 0 to 3-fold change and proteins follow a p-value cutoff of 0.05. Out of scale values are represented in maroon. Non-significant values are left blank and represented in white. **D.** Lollipop chart of biological processes (top panel) and cellular components (bottom panel) enrichment analysis of the secretome dataset obtaind from RPE1 cells treated with siFBL and siNON control for 48 hours, using ShinyGO to select proteins with more than 1.5-fold upregulation and a p-value cutoff of 0.05. Pathways over fold enrichment were plotted with -log10FDR using the indicated color scheme. Number of genes: 44 and FDR cutoff: 0.01. **E.** Heat map representing genes fold change of growth factor, immune response and senescence marker from RNA-seq datasets obtained from RPE1 cells treated with siFBL, siUTP18 and siNON control for 48-, 96- and 192 hours. The scale bar ranges from 0 to 4-fold change and genes follow a p-value cutoff of 0.05. Out of scale values are represented in maroon. Non-significant values are left blank and represented in white. The gene list is based on available literature. **F.** Heat map representing genes fold change of the YAP/TAZ-TEAD signaling pathway from the RNA-seq dataset obtained from RPE1 cells treated with siFBL, siUTP18 and siNON control for 48-, 96- and 192 hours. The scale bar ranges from 0 to 4-fold change and genes follow a p-value cutoff of 0.05. Out of scale values are represented in maroon. Non-significant values are left blank and represented in white. The gene list is based on available literature. **G.** Heat map representing genes fold change of the WNT signaling pathway from the RNA-seq dataset obtained from RPE1 cells treated with siFBL, siUTP18 and siNON for 48-, 96- and 192 hours. The scale bar ranges from 0 to 4-fold change and genes follow a p-value cutoff of 0.05. Out of scale values are represented in maroon. Non-significant values are left blank and represented in white. The gene list is based on available literature. **H.** Representative confocal microscopic images of RPE1 cells treated with the indicated single or double siRNAs for 96 hours (siNON, siFBL, siNON+siTP53, siFBL+siTP53, siUBF, siUTP18, siUBF+siTP53, siUTP18+siTP53), and co-stained for actin stress fibers using phalloidin (red), tubulin TUBB (green) and DAPI (blue). Images are representative of N=4 biological replicates. Scale bar: 20 µm. **I.** Representative confocal microscopic images of RPE1 cells treated with the indicated single or double siRNAs for 96 hours (siUBF, siUTP18, siUBF+siTP53, siUTP18+siTP53), and co-stained for actin stress fibers using phalloidin (red) and DAPI (blue). Corresponding siFBL and siNON control are shown in Fig 7I. Images are representative of N=4 biological replicates. Scale bar: 20 µm. **J.** Representative confocal microscopic images of RPE1 cells treated with the indicated single or double siRNAs for 96 hours (siUBF, siUTP18, siUBF+siTP53, siUTP18+siTP53), and co-stained for actin stress fibers using phalloidin (red) and DAPI (blue). Corresponding siFBL and siNON control are shown in Fig 7I. Images are representative of N=4 biological replicates. Scale bar: 20 µm. **K.** Bar graph showing speed of cells derived from live-cell imaging of FUCCI-RPE1 cells treated with the indicated siRNA (siNON, siFBL, siTP53, siCDKN1A, siFBL + siTP53, siFBL + siCDKN1A) for 0 to 48 hours, starting from siRNA addition. 10 cells per condition per replicate are chosen at the 0-hour time point and followed together with their progeny for continuous tracking of cell mobility. The mean with SD is plotted for N=3 biological replicates. Asterisks denote p values (****, <0.0001; *, <0.05) as calculated by an Ordinary one-way Anova test with Tukey’s multiple comparison. ns, non-significant. **L.** Bar graph showing of the migration speed derived from live-cell imaging of FUCCI-RPE1 cells in the G0/G1 cell-cycle phase treated with the indicated siRNA (siNON, siFBL, siTP53, siCDKN1A, siFBL + siTP53, siFBL + siCDKN1A) for 0 to 48-hours. Each data point represents the average speed of five starting cells and their progeny imaged in single imaging frames. The mean with SD is plotted for N=3 biological replicates. Asterisks denote p value (**, <0.01) as calculated by an Ordinary one-way Anova test with Tukey’s multiple comparison. ns, non-significant.

## Resource Availability

### Lead Contact

Further information and requests for resources and reagents should be directed to and will be fulfilled by lead contact Matthias Peter.

### Materials Availability

All reagents generated in this study can be requested without restriction upon an agreement with a material transfer agreement (MTA).

### Data and Code Availability

The RNA-Seq data have been deposited to the NCBI GEO and are available under accession number GEO: GSE305361.

The mass spectrometry proteomics data have been deposited to the ProteomeXchange Consortium via the PRIDE partner repository with the data set identifier PXD067133.

### Experimental model and subject details Cell culture

Human hTERT-immortalized retinal pigment epithelial cells – hTERT RPE1 (purchased from ATCC), hTERT RPE1 FUCCI cells (Kind gift from Randall W. King, Harvard Medical School) and dCas9-RPE1 cells (Kind gift from Corn Lab, IMHS, ETH Zurich) were cultured in DMEM high glucose GlutaMAX medium (Gibco 31966-021) supplemented with 10% FBS, and 1x Penicillin Streptomycin Glutamine (Gibco 10378-016). Cells were kept at 37°C in 5% CO_2_ incubator. All cell lines were routinely checked for mycoplasma contamination by PCR.

## Method Details

### siRNA and siRNA transfections

ON-TARGETplus siRNA – SMARTpool were purchased from Dharmacon, which were validated by the manufacturer. Cells were plated at 20-25% confluency 24 hours prior to siRNA transfections in 6-well plates. Cells were transfected at 40-50% confluency using Lipofectamine RNAiMAX Transfection Reagent as per the manufacturer’s instructions (Invitrogen, 13778075). SMARTpool siRNAs reagent from Dharmacon were used at a concentration 30 pmol per well of 6 well plate, 100 pmol per 100 mm plate and 200 pmol per 150mm plate. Cells were split in fresh medium at 24 hours post transfection and replated according to the experimental requirements.

### Chemical treatments and antibodies

Treatment with Leptomycin B solution from Streptomyces sp. (Sigma Aldrich, L2913) was performed at a final concentration of 25nM for 2 hours in 96 hours siRNA transfected cells. Similarly, Cycloheximide (Sigma Aldrich, C7698) and Actinomycin D (Sigma Aldrich, A9415) were used at final concentrations of 30 μg/ml and 625nM, respectively, for 2 hours in 96 hours siRNA transfected cells. All primary antibodies were commercially purchased, and validation provided by manufacturer’s data available on their websites. Actin staining was performed using Rhodamine Phalloidin labelling probes (Invitrogen, R415).

### MTT Assay

24 hours post transfections; 10,000 cells were plated in 96-well plates with four technical replicates per condition. The CellTiter Non-Radioactive Cell Proliferation Assay (MTT) assay kit was used as per the manufacturer’s instructions (Promega, G4000). The dye solution was mixed with complete media and incubated for 2 hours at 37°C in a 5% CO_2_ incubator. The reaction was stopped using Stop Mix provided in the kit. Plates were measured on a Molecular Devices microplate reader Versa max at a wavelength of 570nm using SoftMax Pro 7.3.1 software. The relative cell proliferation was calculated by subtracting blank values and normalizing to day 0 and control (siNON) conditions.

### Live cell imaging with RPE1 FUCCI cells

RPE1 FUCCI cells were plated in 6 well glass bottom plates (Cellvis, P06-1.5H-N). 24 hours post plating, siRNA transfection reagents were added, and live-cell imaging was immediately started. Cells were imaged at every 30-minute interval for a period of 72 or 96 hours with 6 positions per well. Images were acquired using inverted wide-field microscope Nikon Ti2 eclipse microscope at 37°C in 5% CO_2_ with humidity. The objective S Fluor 20x NA 0.75 DIC N2 WD 1.0 mm and Orca Fusion BT camera were used. The cells were imaged using GFP (excitation 488 nm, emission 515 nm; GFP-like filter set) and mCherry (excitation 578 nm, emission 641 nm; mCherry-like filter set). The number of cells at different cell cycle phases was quantified using Cell Profiler Software (version 4.2.8) with standard scripts.

### Fixed cell imaging using Immunofluorescence

RPE1 cells were plated 24-hour post transfection in 8-well on cover glass II (Sarstedt 94.6190.802) chambers. At indicated time points – 48 or 96-hour post transfection, the cells were treated with 4% formaldehyde fixative solution (Invitrogen, FB002) for 15 min at room temperature post treatments and washed three times with 1xPBS. Cells were then fixed with 0.4% Triton X-100 in 1xPBS solution for 15 min at room temperature post treatments and washed three times with 1xPBS. Blocking buffer (5% BSA in 1xPBS) was added and cells were incubated for two hours at room temperature. Without washing, cells were proceeded with primary antibody prepared in blocking solution and incubated at 4°C for 12 to 16 hours. Cells were washed three times for 15 minutes each with 1xPBS and incubated with secondary antibody prepared in blocking solution for two hours at room temperature. All primary and secondary antibodies were used at 1:100 dilution in blocking solution. Cells were washed three times for 15 minutes each with 1xPBS. DAPI was added for 30 min at room temperature in 1xPBS and images were acquired using a Zeiss LSM880 confocal microscope with a 63x oil immersion objective lens at a zoom range 0.6-2x. Z stacks were recorded and collapsed into a single maximum intensity projection using Zen black software. Images were processed using Zen blue or ImageJ software. Images were taken using identical intensity adjustments per experimental setup.

### Transmission Electron Microscopy

RPE1 cells were plated in 150 mm plates for TEM. Cells transfected for 96 hours with siRNA were harvested using disassociation reagent Trypsin-EDTA 0.25% (Gibco, 25200056) and washed with PBS. The samples were initially fixed in 4% paraformaldehyde in PBS for one hour, washed with PBS and then processed using the NAMA-Ur technique^106^. Samples were immersed in 4% paraformaldehyde in 0.5M NaOH overnight at 4°C. The samples were then washed with distilled water, immersed in 1% acetic acid, and washed again in distilled water. Dehydration was accomplished using a graded serious of methanol (25%, 50%, 75%) followed by an acetic anhydride and methanol 5:1 mixture and left to stand overnight in darkness at 25°C, stirred occasionally. Samples were further dehydrated in 100% methanol, followed by 100% ethanol, and then then infiltrated in a graded ethanol-Epon resin (Electron Microscopy Sciences, Hatfield, PA) series at 30%, 50%, 70% and finally at 100% Epon resin. Polymerization of Epon resin was achieved at 60 °C for 72 hr. Ultrathin sections of 60 nm were obtained with a diamond knife (Diatome Ltd., CH) using a Leica UC7 ultramicrotome (Leica Microsystems, CH), placed on Formvar/carbon coated TEM grids (Quantifoil, DE), and stained with 5% aqueous uranyl acetate for 60 minutes at room temperature, followed by 20 minutes at 60°C. Micrographs of the stained sections were imaged using the Thermo Fisher Scientific (TFS) Talos L120C TEM (Thermo Fisher Scientific, USA) equipped with a CMOS camera BM-Ceta, operating at 120 kV acceleration voltage in bright field mode.

### Quantitative PCR with reverse transcription

RPE1 cells were harvested 24 hours or 96 hours post siRNA transfections in TRIzol reagent (Ambion Life tech, 15596018) and RNA was extracted as per manufacturer’s instructions. The extracted RNA was subjected to genomic DNA elimination reaction and reverse-transcription assay using Takara PrimeScript RT reagent kit with gDNA Eraser (Takara, RR047A) as per manufacturer’s guidelines. The qPCRs were done using MasterMix Roche KAPA SYBR FAST reagent (KAPA Biosystems, KK4611) and a Roche Light Cycler 480 was used for measurements. DNA amounts were quantified using the ΔΔCt method, and the control (siNON) condition was set to 1. Actin was used to normalize the assay.

### Polysome Profiling

RPE1 cells were harvested 96 hours post siRNA transfections and washed once with 1xPBS containing 100μg/mL CHX. 10% and 50% sucrose solutions were made in 1x sucrose buffer (50mM HEPES pH7.5, 100mM KCl, 10mM MgCl_2_). Lysis buffer (10mM Tris pH7.5, 100mM KCl, 10mM MgCl_2_, 1% Triton X-100, 1mM DTT, 100μg/mL CHX, 1X Protease inhibitors, 1mM PMSF) was added to the cells and incubated at 4°C rotating for 20 minutes. Cell lysates were precleared at 1000g for 10 minutes at 4°C. Protein concentration was measured using BCA and equal amounts were loaded on sucrose cushions. The samples were ultracentrifuged at 39000 rpm for 2 hours at 4°C using Optima XPN-100 and a SW41 rotor. Absorbance of the fractions was measure at 254nm. The sucrose gradients and polysome profile measurements were made using TRIAX fractionator from BIOCOMP Instruments.

### Nascent transcription and translation assays

Cells were incubated with 1mM 5-Ethynyl-uridine (5-EU, Jena Bioscience, CLK-N002-10) for 1 hour and 3 hours for labelling nascent RNA and 10μM O-Propargyl-puromycin (OPP, Jena Bioscience, NU-931-05) for 3 hours for nascent protein labelling. The click chemistry reaction was performed using CuAAC Cell Reaction Buffer Kit (BTTAA based) as per manufacturer’s instructions (Jena Bioscience, CLK-073). Briefly, the cells were fixed, permeabilized and incubated with blocking solutions using a fixed cell imaging protocol. 200μl of CLICK reaction buffer containing 20μM AF488-Azide (Jena Bioscience, CLK-1275-1) was incubated for 60 minutes. The CLICK reaction buffer was removed, and the cells washed three times with 1xPBS. DAPI was added for 30 minutes at room temperature in 1xPBS and images were acquired with a Zeiss LSM880 confocal microscope using 63x oil immersion objective at 0.6x zoom range. Z stacks were recorded and collapsed into a single maximum intensity projection using Zen black software. Images were processed using Zen blue and ImageJ software. **Signal Quantification:** To quantify signal intensities, standard measurement features in FIJI 1 software were used. Average signal intensities were recorded from small region of interest (ROI) in the nucleolus, nucleus and cytoplasm, respectively, avoiding void spaces. The raw intensity values were normalized to average of values from control (siNON) adjusted to 1 separately for every individual experiment performed. The data was plotted, and statistics was performed using GraphPad PRISM 10.

### Western Blotting

Whole cell lysates were prepared in lysis buffer (50mM HEPES pH 7.5,150mM NaCl, 1% NP-40, 1% Triton X-100, 0.1% SDS, 7% glycerol, 1X Protease inhibitors, 1mM PMSF). Samples were mixed in 3:1 ratio with 4x protein loading buffer (200 mM Tris-HCl pH 6.8, 8% SDS, 40% glycerol, 0.2% bromophenol blue and 20% beta-mercaptoethanol) and boiled for 10 minutes at 95°C. Bolt 4-12% Bis-Tris gels (Invitrogen, NW04127) in MOPS SDS buffer was used to run samples followed by standard western blot procedures with wet transfer system using 0.45 μ m nitrocellulose blotting membrane (Amersham Protran, 10600002). Membranes were blocked in 5% milk in 1xPBST (0.01% Tween 20). Primary antibody were diluted in blocking solutions ranging between 1:100 to 1:1000 and incubated overnight at 4°C. Secondary antibody was diluted in blocking solution ranging between 1:1000 to 1:10000 with 2 hours incubation at room temperature. Blots were developed with SuperSignal West Pico PLUS or Femto solutions (Thermo Fisher, 1859022) and scanned on a Fusion FX7 imaging system (Witec AG).

### RNA extraction and next generation sequencing

#### RNA Extraction

RPE1 cells were harvested from 6-well plates at 48, 96 and 192 hours post siRNA transfection. Three independent replicates were transfected and harvested simultaneously for each knockdown condition (siNON, siFBL and siUTP18). The cells were washed with 1xPBS, and pellets frozen at -80°C until processing. RNA was purified using RNeasy PlusMini Kits as per manufacturer’s instructions (Qiagen, 74134). RNA concentrations were measured using Nanodrop One (Witec AG).

#### Library preparation

The quality of the isolated RNA was determined with a Fragment Analyzer (Agilent, Santa Clara, California, USA). The TruSeq Stranded Total RNA Library Prep Gold (Illumina, Inc, California, USA) was used in the succeeding steps. Briefly, total RNA samples (100-1000 ng) were depleted with ribosomal RNA and then reverse-transcribed into double-stranded cDNA. The cDNA samples were fragmented, end-repaired and adenylated before ligation of TruSeq adapters containing unique dual indices (UDI) for multiplexing. Fragments containing TruSeq adapters on both ends were selectively enriched with PCR. The quality and quantity of the enriched libraries were validated using a Fragment Analyzer (Agilent, Santa Clara, California, USA). The product was a smear with an average fragment size of approximately 260 bp. The libraries were normalized to 10nM in Tris-Cl 10 mM, pH8.5 with 0.1% Tween 20.”

#### Cluster Generation and Sequencing

The NovaSeqX (Illumina, Inc, California, USA) was used for cluster generation and sequencing according to standard protocol. Sequencing was paired end at 2 X150 bp.“

#### RNA-Seq Data Analysis

The RNA-Seq analysis was performed in several steps. Raw sequencing reads were pre-processed using Fastp (v0.23.4)^107^ to remove adapter sequences, trim low-quality bases, and filter out reads with a Phred quality score below 20.

Read alignment was conducted with STAR (v2.7.10b)^108^ using the Ensembl genome build GRCh38.p13 as reference, with gene annotations from GENCODE release 42 (downloaded on 2023-01-30). The following STAR parameters were used: --sjdbOverhang 150 --outFilterType BySJout --outFilterMatchNmin 30 --outFilterMismatchNma× 10 –outFilterMismatchNoverLmax 0.05 --outMultimapperOrder Random --alignSJDBoverhangMin 1 --alignSJoverhangMin 8 -- alignIntronMa× 100000 --alignMatesGapMa× 100000 --outFilterMultimapNmax 50 -- chimSegmentMin 15 --chimJunctionOverhangMin 15 --chimScoreMin 15 –chimScoreSeparation 10 --outSAMstrandField intronMotif --alignEndsProtrude 3 ConcordantPair.

Gene-level expression was quantified using the feature Counts function from the R subread package (v2.14.1)^109^. The following options were applied: minimum mapping quality of 10, minimum feature overlap of 10 bp, inclusion of multi-mapping reads, counting of only primary alignments, and counting of reads overlapping multiple genes.

Differential gene expressions were assessed using the generalized linear model framework of DESeq2 (v1.40.1)^110^, running on R version 4.3.0. Genes with an adjusted p-value (Benjamini-Hochberg method) below 0.05 were considered differentially expressed.

The PCA and volcano-plots are generated using FGCZ Shiny Apps.

### Mass spectrometry for Protein abundance profiling

#### Protein Extraction

RPE1 cells were harvested from 100 mm plates at 48, 96 and 192 hours post siRNA transfection. Four independent replicates were transfected and harvested simultaneously for each knockdown condition (siNON and siFBL). The cells were washed with 1xPBS, and pellets frozen at -80°C until processing. The cell pellets were resuspended in 8M Urea lysis buffer (8M urea, 50mM Tris HCl pH 7.5, 75mM NaCl, 10mM NaF, 10mM sodium pyrophosphate, 5mM 2-glycerophosphate, protease and phosphatase inhibitor cocktail), sonicated using a Bioruptor at medium amplitude for 5 cycles of 30 seconds ON and 30 seconds OFF, and centrifuged at 13,000 rpm for 15 minutes at room temperature to pellet debris. The protein concentration was measured using a BCA protein quantification assay.

#### Protein digestion

2 mg total protein was transferred in a fresh low bind tube, Tris (2-carboxyethyl) phosphine (TCEP, Sigma-Aldrich, C4706) was added to a final concentration of 5mM and incubated for 30 minutes at room temperature. Iodoacetamide (IAA, Sigma-Aldrich, I1149) was then added to final concentration of 10mM and incubated for 30 minutes at room temperature in the dark. Samples were then diluted to adjust the urea concentration to 4M using dilution buffer (50mM Tris HCl pH 7.5, 75mM NaCl) and incubated with 1:200 LysC to protein ratio for 2 hours shaking at room temperature. Subsequently, the samples were diluted to 2M urea concentration using dilution buffer, trypsin was added at 1:100 trypsin to protein ratio and incubated overnight shaking at 37°C.

#### Column clean-up

Digestion was quenched with 5% formic acid and samples were cleaned up with C18 columns using BioPure SPN MIDI (The Nest Group, HEM S18V) following manufacture indications.

#### Analysis

The samples were acquired on an Orbitrap Exploris 480 mass spectrometer (Thermo Scientific) coupled to a Vanquish Neo UHPLC system (Thermo Scientific). Peptides were separated using a C18 reversed phase column (75 μm x 400 mm (New Objective), packed in-house with ReproSil Gold 120 C18, 1.9 μm (Dr. Maisch GmbH)), using a 120-minute linear gradient from 7% to 35% buffer B at a flow rate of 300 nl/min (buffer A: 0.1% [v/v] formic acid, buffer B: 0.1% [v/v] formic acid, 80% [v/v] acetonitrile). The mass spectrometer was operated in data-dependent acquisition (DDA) mode with the following parameters: one MS1 scan (350-1500 m/z, 60000 resolution, 100% normalized AGC target, auto maximum injection time), followed by a maximum of 20 MS2 scans (1.4 m/z isolation window, 15000 resolution, 200% normalized AGC target, auto maximum injection time). Intensity threshold was set to 5.0e3, charge states 2-6 were included, and a dynamic exclusion of 25 seconds was applied. Ions were fragmented with HCD (NCE 28%).

**Raw MS data** were searched against the UniProt human proteome (Swiss-Prot & TrEMBL, version 2024_03) using MaxQuant (v2.6.1.0) with the Andromeda search engine. Trypsin/P was specified as the enzyme for specific digestion with up to 2 missed cleavages. First search peptide tolerance was set to 20 ppm. Carbamidomethyl (C) was set as fixed, and oxidation (M) and acetyl (N-term) as variable modifications. Label-Free Quantification (LFQ) was used with match between runs disabled. Unique and razor peptides were used for quantification. PSM and Protein FDR were set to 0.01.

MaxQuant output was further processed using Perseus (v2.0.11). Protein groups were filtered for having valid values in all replicates of at least one condition. Missing values were imputed from a normal distribution. Volcano plots were generated by conducting Student’s T-test using a permutation-based FDR of 0.05 and plotting the results in R.

### Mass spectrometry for Secretome

#### Protein Extraction

Supernatant complete media from RPE1 cells were harvested from 100 mm plates at 48- and 96-hours post siRNA transfection or supernatant starvation media without FCS for 24 hours pre-harvesting at 96 hours post siRNA transfection. Three independent replicates were transfected and supernatant media harvested simultaneously for each knockdown condition (siNON and siFBL). The supernatant media was centrifuged at 10,000g at room temperature for 10 minutes. The supernatant was concentrated using Ultracentrifuge filters 3KDa MWCO (Amicon UFC800324) from 5ml to 400μl. Samples were spun at 13,000 rpm for 15 minutes at room temperature to remove debris. The protein concentration in the supernatant was measured using the BCA protein quantification assay. Equal ratios of 8M Urea lysis buffer (8M urea, 50mM Tris HCl pH 7.5, 75mM NaCl, 10mM NaF, 10mM sodium pyrophosphate, 5mM 2-glycerophosphate, protease and phosphatase inhibitor cocktail) was added, and protein reduction, column clean-up, and MS analysis performed as described above.

#### Analysis

LC-MS/MS analysis was performed on an Orbitrap Exploris 480 mass spectrometer (Thermo Scientific) coupled to a Vanquish Neo UHPLC system (Thermo Scientific). Peptides were separated using a C18 reversed phase column (75 μm x 400 mm (New Objective), packed in-house with ReproSil Gold 120 C18, 1.9 μm (Dr. Maisch GmbH)), using a 60-minute non-linear gradient from 1% to 43.7% buffer B at a flow rate of 300 nl/min (buffer A: 0.1% [v/v] formic acid, buffer B: 0.1% [v/v] formic acid, 80% [v/v] acetonitrile).

The mass spectrometer was operated in data-independent acquisition (DIA) mode with the following parameters: one MS1 scan (330-1650 m/z, 120000 resolution, 300% normalized AGC target, 20 ms maximum injection time), followed by 30 variable MS2 windows from 330 to 1650 m/z with 1 m/z overlap (30000 resolution, 2000% normalized AGC target, 64 ms maximum injection time). Ions were fragmented with HCD (NCE 27%).

**Raw MS data** were searched against the UniProt human proteome (Swiss-Prot & TrEMBL, version 2024_03) using Spectronaut (v19, Biognosys AG) in directDIA mode. Trypsin/P was specified as the enzyme for specific digestion with up to 2 missed cleavages. Carbamidomethyl (C) was set as fixed, and oxidation (M) and acetyl (N-term) as variable modifications. PSM, Peptide, and Protein Group FDR were set to 0.01. MS1 and MS2 mass tolerance strategies were set to +/-40 ppm. Quantification calculations were performed on the MS2 level using the XIC area and cross-run normalization was enabled.

### Gene Ontology (GO) Enrichment Analysis

Gene set enrichment analysis was performed with the ShinyGO 0.82 web browser (https://bioinformatics.sdstate.edu/go/), using the parameters mentioned in the corresponding figure legends^111^. Heat maps are generated using PRISM version 10 with genes and proteins based on limited literature review.

### Genome-wide CRISPRi screen

dCAS9-KRAB RPE1 cells were transduced with the pCRISPRia-v2 lentiviral library containing 100,000 sgRNAS at a multiplicity of infection (MOI) of approximately 0.2 to 0.3 with a coverage of at least 300 cells per sgRNA in a media supplemented with 4 µg/ml protamine sulphate. The media was changed 24 hours post transduction, containing 15 µg/ml puromycin. The selection was maintained for 96 hours, and MOI was calculated to be 0.23. The cells were expanded for another 7 days and then divided into two parts. The first half was siRNA transfected with control siRNA (siNON) and the second half with fibrillarin siRNA (siFBL). The media was changed at 24 hours and cells were regularly split every 2-3 days. The samples were collected after 5, 10 and 15 days (120, 240 and 360 hours) post siRNA transfection and cell pellets frozen at -80°C until processing. Genomic DNA was isolated using a Purelink Genomic DNA isolation kit (Invitrogen, K182001) as per manufacturer’s instructions.

Genomic DNA was quantified using NanoDrop® ND-1000 UV-Vis Spectrophotometer. To reach the theoretical coverage after sequencing 10 µg of gDNA was amplified in parallel one-step PCRs with barcoded primers targeting the pCRISPRia-v2 vector backbone. The primers used were CRISPRi_i5-1 and CRISPRi_i7-1 on the the first PCR followed by the NEBNext® Multiplex Oligos for Illumina® (96 Unique Dual Index Primer) on the second PCR. Primers were ordered as desalted Oligos at IDT (Integrated DNA Technologies, LubioScience GmbH). PCR conditions were as follows: 10µg of gDNA per 100 µl reaction, x units of Q5 polymerase (New England Biosciences), and 23-27 cycles of amplification.

The amplified product containing the spacer sequence with a size of around 300 bp was purified by SPRI bead cleanup and a right-sided size selection to remove the genomic DNA input. Using a dual barcoding system, the final NGS amplicons were multiplexed and sequenced on Illumina platform NextSeq 2000 with 21 dark cycles (DC) in Read 1 to skip the invariable region of the primer binding side (U6 promoter). Sequencing run parameters: 21 DC + 36 bases R1, 8 bases I1, 8 bases I2 and a spike in of 5-10% of PhiX Control v3 (Illumina). Loading concentration on a NextSeq 2000 is 400pM. Bcl files were demultiplexed using bcl2fastq (Illumina) and allowing 0 mismatches in Indexes 1 and 1 mismatch in Index 2. For a data quality check we ran FastQC (Babraham Bioinformatics) to display the base quality scores and an inhouse pipeline mapping a subset of the sgRNA read outs against commonly used CRISPR libraries (R package ezRun (https://github.com/uzh/ezRun) through the data analysis framework SUSHI^112^.

### CRISPR Screen Analysis

For CRISPR screen data, raw reads were trimmed at the 3′ end by 17 bases using Fastp to extract the sgRNA sequences. Trimmed reads were aligned to the CRISPRi human reference library using Bowtie, allowing up to one mismatch.

Read counts were aggregated at the gene level, and differential representation across conditions was evaluated using the edgeR package, which models count data using negative binomial distributions

### Quantitative phase imaging (QPI)

RPE1 cells were plated in 6 well glass bottom plates (Cellvis, P06-1.5H-N). 24 hours post plating, siRNA transfection reagents were added, and live-cell imaging was immediately started using a Holotomographic quantitative phase imaging microscope (Tomocube X1) at 37°C in 5% CO_2_ with humidity. Cells were imaged at 30-minute intervals for a period of 72 or 96 hours. For fixed samples, RPE1 cells were plated in 6 well glass bottom plates (Cellvis, P06-1.5H-N), and 24 hours post plating, siRNA transfection reagents were added. Cells were fixed at 96 hours post siRNA transfection using 4% formaldehyde fixative solution (Invitrogen, FB002) for 15 min at room temperature post treatments and washed three times with 1xPBS. The refractive index (RI) of nucleoli in each cell was measured by using an ImageJ script that projects tomographic images and expands one pixel outward from the brightest point and calculates the median value.

### Cell tracking pipelines

RPE1 FUCCI cells were imaged for live-cell analysis, and cell tracking was performed using the MATLAB plugin CellMAPtracer (Ghannoum et al., 2021), which enables automated tracking and cell cycle phase prediction for FUCCI-labelled cells. Automated tracks were manually curated to correct tracking errors and to annotate cell divisions. Phase prediction outputs were visually inspected and corrected as needed due to occasional inaccuracies caused by imprecise centroid localization. For quantification of cell speed across treatments, average speed values generated by CellMAPtracer were used. Five cells per imaging position were tracked over time, with six positions analyzed during the 48 to 96-hour window and two positions during the 0–48 hour interval. To evaluate how cell speed varies with cell cycle phase, instantaneous speeds were manually calculated from positional changes of cell centroids, as these values were not directly provided by CellMAPtracer. Speed between consecutive frames was computed using the formula: Speed=(sqrt(X_t+1_−X_t_)^2^+(Y_t+1_−Y_t_)^2^)x0.32 μm)/0.5 hour where 0.32 µm corresponds to the spatial calibration (1 pixel = 0.32 µm), and 0.5 h is the imaging interval (30 minutes). Speeds were averaged over five cells per position, including daughter cells, across the entire 48-hour imaging period, separately for G1 and combined S/G2/M phases based on CellMAPtracer phase predictions.

